# Projective LDDMM: Mapping Molecular Digital Pathology with Tissue MRI

**DOI:** 10.1101/2022.04.22.489163

**Authors:** Kaitlin M. Stouffer, Menno P. Witter, Daniel J. Tward, Michael I. Miller

**Affiliations:** Department of Biomedical Engineering, Johns Hopkins University, Baltimore, MD, USA; Kavli Institute for Systems Neuroscience, Norwegian University of Science and Technology, Trondheim, Torgarden, Norway; Departments of Computational Medicine and Neurology, University of California, Los Angeles, CA, USA

**Author notes:** Contributing authors. These authors contributed equally to this work (Daniel J. Tward, Michael I. Miller).

**Keywords:** Projective LDDMM, Multimodal and Multiscale Image Registration, Digital Pathology

## Abstract

Reconstructing dense 3D anatomical coordinates from 2D projective measurements has become a central problem in digital pathology for both animal models and human studies. We describe a new family of diffeomorphic mapping technologies called Projective LDDMM which generate diffeomorphic mappings of dense human MRI atlases at tissue scales onto sparse measurements at micron scales associated with histological and more general optical imaging modalities. We solve the problem of dense mapping surjectively onto histological sections by incorporating new technologies for crossing modalities that use non-linear scattering transforms to represent multiple radiomic-like textures at micron scales and incorporating a Gaussian mixture-model framework for modelling tears and distortions associated to each section. We highlight the significance of our method through incorporation of neuropathological measures and MRI, as relevant to the development of biomarkers for Alzheimer’s disease and one instance of the integration of imaging data across the scales of clinical imaging and digital pathology.

## 1 Introduction

The past decade has ushered an “omics” revolution into biomedical research, with high yields of data ranging from microscopic to macroscopic scales. Modern machine learning methods coupled with image processing have enabled the integration of “pathomics” data extracted from digital pathology technologies with “radiomics” data extracted from lower resolution imaging technologies such as magnetic resonance imaging (MRI) in a number of niche applications, such as those within the domain of cancer diagnostics and prognostics [1, 2]. However, approaches remain widely varied across applications and often require particularities in image acquisition and image type, such as block face imaging [3], to facilitate alignment between imaging modalities [4]. The obstacle of registering spatially incomplete sets of 2D (or 3D) images to a dense 3D atlas remains a particular challenge in both animal and human settings, with no approach yet amenable to synthesizing the imaging data from the spectrum of technologies within this domain.

This paper introduces a new class of image-based diffeomorphometry methods which we term Projective LDDMM for aligning sparse sets of image captures to 3D coordinate systems across micron and millimeter scales. We focus particularly on the registration of 3D MRI with 2D digital histology, as representative of the class of multi-scale, multi-modality mapping in biomedical research including traditional light microscopy mapping to dense reference atlases [5, 6, 7, 8], light sheet methods [9, 10], deep tissue imaging [11], and spatial transcriptomics [12, 13, 14, 15]. We formulate the mapping of dense atlases to sparse images problem using the random orbit model of computational anatomy [16, 17, 18, 19] in which the space of dense anatomies 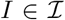 is modeled as an orbit of a 3D template under the group of diffeomorphisms. Projective LDDMM models the sparse 2D observables not as an element of the orbit 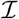 but rather a random deformation in dense 3D coordinates composed with a measurement projection to sparse coordinates. This implies that the random orbit model encompasses the composition of two observation channels: one for projection and one for post-projection processing, such as the steps involved with histological staining and slide preparation. While LDDMM provides the geodesic metric [20, 21] on the orbit of 3D anatomies, there is no symmetry between the observable and the template, in general, and there should not be. This departs significantly from the symmetric methods [22, 23].

Alignment specifically of modes of histology to MRI warrants two extensions of the basic model of Projective LDDMM. First, cross-modality similarity modelling is essential. Several strategies for representing image similarity have emerged including cross-correlation [24], mutual information [25], and local textural characteristics [26]. Our approach extends previous work [3, 27] modelling a photometric transformation of histology to MRI by expanding histology image space to a span of discriminative “filtered images” via Mallat’s Scattering Transform [28, 29]. These “filtered images” represent local radiomic textures at histological scales and are used to predict MRI contrast. Second, histological images carry large numbers of imperfections with tears, image stitching, and lighting variations. Extending previous work [6, 27], we introduce Gaussian mixtures models in the image plane of each histological slice to interpret image locations as matching tissue, background, or artifact. We proceed by way of the Expectation-Maximization (EM) algorithm [30] in estimating deformations that prioritize image matching at locations that are, in turn, estimated more likely to be matching tissue.

Here, we use Projective LDDMM to reconstruct the 3D geometry of 2D histological sections taken from the medial temporal lobe (MTL) of a brain with advanced Alzheimer’s disease (AD). Efforts to diagnose and manage AD earlier in its disease course have centered on the identification of biomarkers [31]–measures shown to correlate with disease course, but without establishment in AD’s pathological underpinnings of misfolded proteins (tau tangles and amyloid-beta (A*β*) plaques [32, 33, 34]. As one application of our method, the reconstruction of a complete spatial profile of tau pathology at the micron level is necessary for validating such biomarkers as entorhinal cortex thinning [35]. Following registration, we extract quantities of tau pathology using a machine learning-based approach that are then mapped to 3D via the correspondences yielded by Projective LDDMM. We demonstrate the efficacy of modeling these quantities using a measure-based framework [36] as befits resampling the quantities at different scales and within both 2D manifolds and 3D volumes for correlation with biomarkers of interest.

## 2 Results

### 2.1 Projective LDDMM

In the random orbit model of computational anatomy [16], the unobserved space of human anatomical images, *I*: *R*^3^ → *R^r^*, is modeled as an orbit under diffeomorphisms of a template

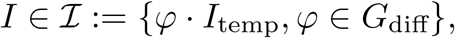

*G*_diff_ the group of diffeomorphisms *φ*: *R*^3^ → *R*^3^, which act on images as *φ* · *I* = *I* ○ *φ*^-1^. The observables *J*: *R*^3^ → *R^q^* are modelled as a random field with mean due to the randomness of diffeomorphic deformation and measurement process. For different problems of interest, the atlas image is *R^r^*-valued with, for instance, *r* = 1 corresponding to single contrast MRI or *r* = 6 for diffusion tensor images (DTI) [37]. Likewise, observables are 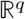-valued with *q* = 3 for traditional histological stains corresponding to the red, green, and blue channels, or *q* >> 6 for alternative representations, such as that given by the Scattering Transform [28] encoding meso-scale radiomic textures in histology images (see Section 4.2). In general, the range space of 3D templates versus targets do not have the same dimension, so *q* ≠ *r*.

Projective LDDMM is characterized by the fact that the observable is not dense in the 3D metric of the brain. Rather, the observable(s) result from either optical or physical sectioning, as in histological slice preparation, taking LDDMM into the projective setting akin to classical tomography [38, 39]. This sectioning lends itself to extending the random orbit model of computational anatomy involving a single channel of Shannon theory to include a second channel, which represents the projective geometry. As shown in Figure 1, this projection channel precedes a second channel associated to the parameters of processing post sectioning, such as rigid motions and diffeomorphic deformations.

**Fig. 1.**
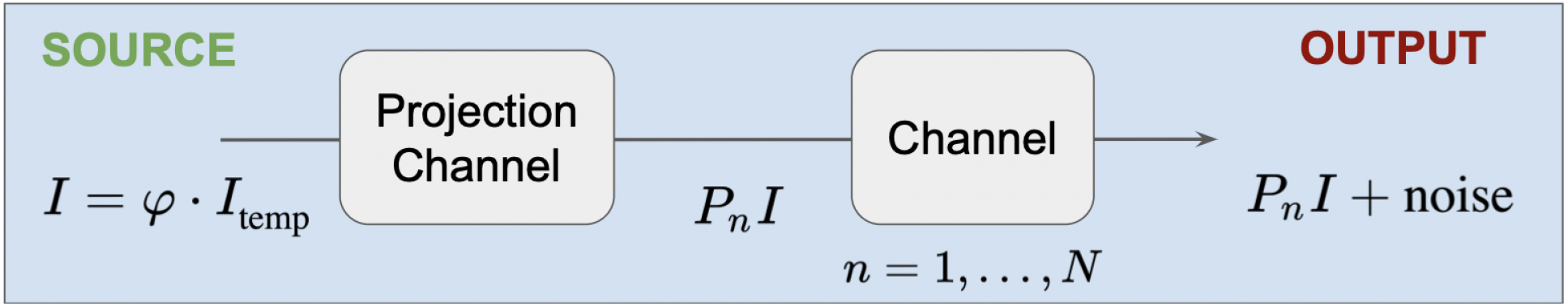
The random orbit model, *φ*·*I*_temp_, extended with projection operator *P_n_*, to generate a set of noisy observables *P_n_I*+noise, *n* = 1, …, *N*, representative of histological processing of tissue post slicing.

The sample measured observables *J_n_*(·), *n* = 1,2, … are a series of (transformed) projections *P_n_* of *I*(·) on the source space *X* ⊂ *R*^3^ to measurement space *Y* ⊂ *R*^2^ (or *Z* ⊂ *R*^1^) defined through the class of point-spread functions associating source to target:

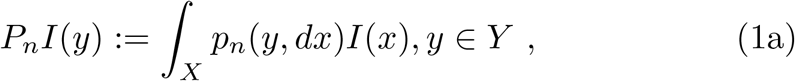

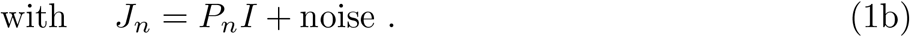

We adopt measure theoretic notation, *p_n_*(*y, dx*) for describing point-spreads to accommodate those taking the form of generalized functions, such as the delta Dirac. Density notation *δ*(*x* – *x*_0_)*dx* corresponds to the measure notation *δ*_*x*o_(*dx*). Each evaluated against a test function *f* (*x*) ∈ *C*^0^ yields *f* (*x*_0_).

The diffeomorphism *φ* is generated as the solution to the flow

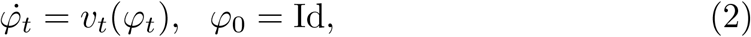

where Id is the identity map and with velocity field *ν_t_*, *t* ∈ [0,1] controlling the flow constrained to be an element of a smooth reproducing kernel Hilbert space (RKHS) 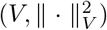 with the entire path square integrable 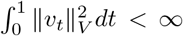 ensuring smoothness and existence of the inverse [40].

This gives us the first variational problem of Projective LDDMM, with || · ||_2_ defined to be the L2 norm.

#### Variational Problem 1 (Projective LDDMM)

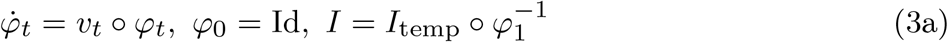

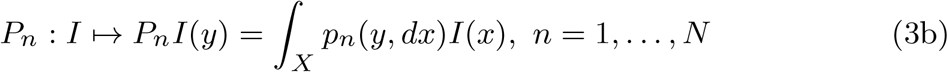

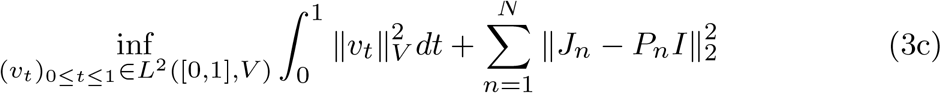

The model specific to our histology images projects the volume *I*(·), defined as a function with domain *X* ⊂ *R*^3^, to parallel sections *J_n_*(·) on *Y* ⊂ *R*^2^, along the third (*z*) dimension, with coordinates, *z_n_, n* = 1, ⋯, *N*. This represents a surjection from source space *X* to target space *Y*, where regions of the 3D volume remain unmapped to the set of target sections. For this, we define ‘Dirac’ point-spreads from *δ_x_* applying to infinitesimal volumes in space (*dx*) with *δ_x_*(*dx*) equal to 1 if *x* ∈ *dx*, and 0, otherwise. Our Dirac point-spreads *p_n_*(*y,dx*) = *δ_y,Z_n__*(*dx*) with (*y, z_n_*) = (*y*^(1)^, *y*^(2)^, *z_n_*) ∈ *R*^3^ concentrate on the planes:

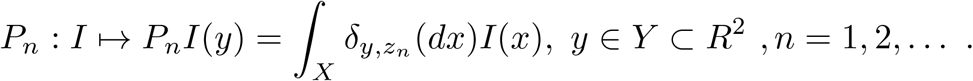

In the case that source and target space are of the same dimensionality and our projection operator *P_n_* is the identity, (3c) reduces to the classic variational problem associated to the random orbit model [16]. In the following section, we introduce an additional complexity of expanding images in a basis for crossing modalities of histology and MRI. While we choose to expand our target images, a similar expansion of the template image yields a span of possible templates, achieving the setting of multi-atlas models popularly used [41, 42, 43].

### 2.2 Projective LDDMM with Scattering Transforms for Crossing Modality and Contrasts in Digital Pathology

In digital pathology, different imaging contrasts emerge from the variety of stains used to elucidate different molecules, such as myelin (LFB), amyloid (6E10), and tau (PHF-1). For crossing from the range space of any of these histology contrasts to MRI, we define a predictive basis, 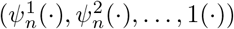, using PCA on a set of nonlinear basis images generated from our observables, *J_n_*(·), *n* = 1, …, *N*. This process is summarized in Figure 2. We expand each histology image to a basis of “filtered images”, each a different scale and texture determined by Mallat’s Scattering Transform [28, 29] through alternating wavelet convolutions and nonlinear modulus operators across scales (see Supplementary Note S.3), followed by PCA reduction:

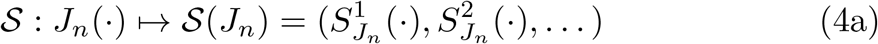

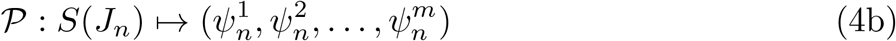

**Fig. 2.**
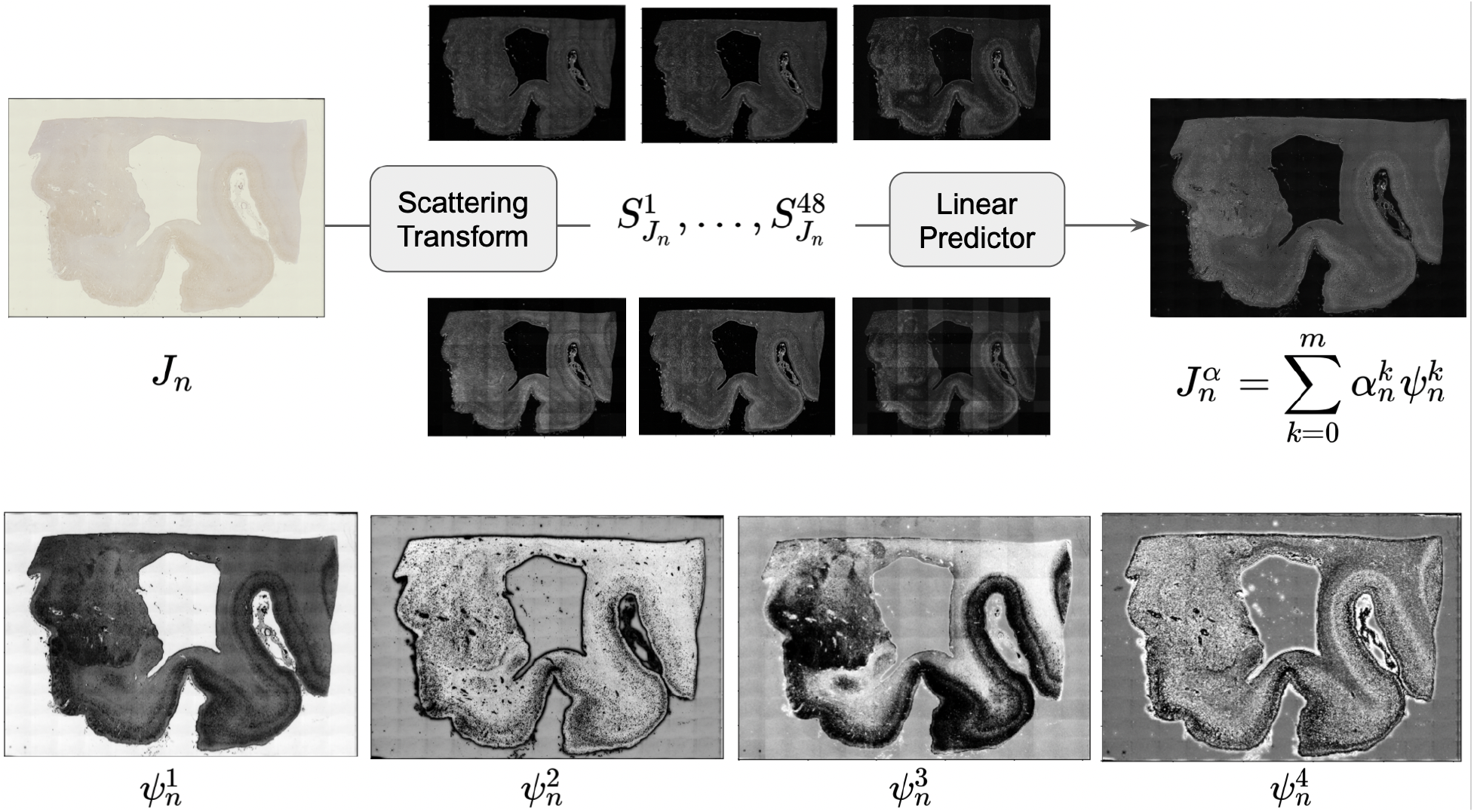
Initial histology image *J_n_* at 2 *μ*m resolution (left). Scattering images generated via a Scattering Transform (middle). Output of linear predictor, 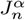, predicting MRI contrast from Scattering images (right). Scattering images projected onto first 4 elements of 6-dimensional PCA basis (bottom).

From the nonlinear filtered images of (4a), we generate the predictive functions, 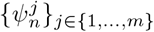, using PCA and adding a constant image, 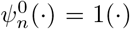. From this basis, we then predict the MRI contrast of our transformed and projected observable as a least squares minimization, written as 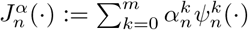.

For digital pathology and other surjective measurements, there are additional parameters associated with deformation of tissue section geometry independently in each imaging plane. We denote the associated rigid and/or affine motion in each imaging plane, *ϕ_n_* ∈ Φ: *R*^2^ → *R*^2^. Estimation of *α_n_* and *ϕ_n_* together with *φ* gives Variational Problem 2.

#### Variational Problem 2

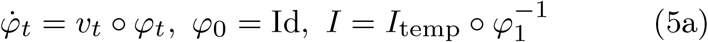

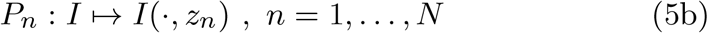

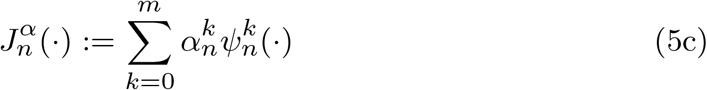

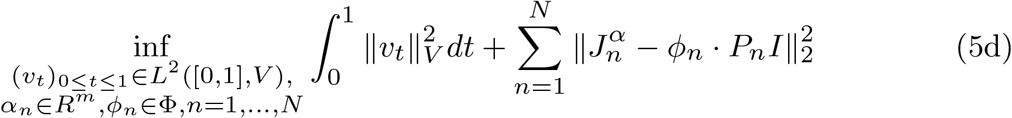

For rigid and/or affine motions, as used in modeling block sectioning in [5], we apply *ϕ_n_* to the histology images and estimate these deformations alternately with deformations of the template (see Algorithm 1). Herein, we expand our dimensions to non-rigid deformations *ϕ_n_* ∈ Φ the group of diffeomorphisms on *R*^2^. A penalty term is introduced 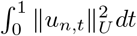 to (5d), with 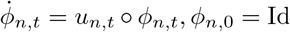. For estimating the cross-modality dimensions *α_n_*, we treat them in a maximum-likelihood setting, optimizing them with initial conditions of deformation and image plane dimensions fixed, then solve the variational problem over all of the other dimensions with the *α_n_* estimates fixed (see Section 4.2). This avoids collapse of the variational problem in these high dimensional settings.

The independent processing of sections requires introduction of additional models for interpreting the measurements in each histology tissue section modelling the tissue as foreground, artifacts of tears and distortion, and background (see Tward et. al [27]). The histology is modelled as a conditionally Gaussian mixture model with means 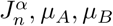 representing foreground tissue, artifacts, and background:

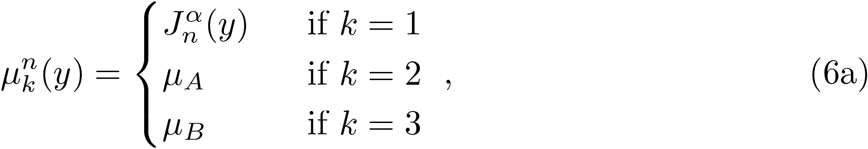

with norm-square term in (5d) replaced by

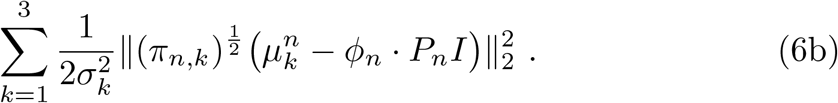

Weighted least-squares interprets the images weighing each model *π_n,k_*(·) with 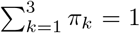 and 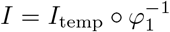 as above. The weights are estimated iteratively, arising from the E-step of an Expectation-Maximization (EM) algorithm [30] selecting at each point in the image the appropriate model for giving the spatial field of weights. This iteration corresponds to a Generalized EM (GEM) algorithm [30] (see Section 4.4 for proof). The results highlighted in Section 2.4 were generated following the approach of this section.

### 2.3 Optical Sectioning, PET, and Parallel Beam Tomography

Additional modes of imaging introduce settings of ideal and non-ideal planar and linear projections fitting the framework for Projective LDDMM. Confocal optical sectioning reconstruct volumes *X* ⊂ *R*^3^ with models that are fundamentally 3-D point-spreads *p_n_*(*y, dx*), *y* ∈ *R*^3^, *dx* ⊂ *R*^3^ with uncertainty supported over the volumes [44, 45], with imaging focused to *n* = 1, …, *N* measurement planes with significant blur out of plane. The mean field of the measurement volume *J_n_*(·) on *R*^3^ are given by the projections *P_n_I*(·) on *R*^3^ of (3b).

Two-dimensional (2D) positron emission tomography (PET) introduces point-spreads *p_n_*(*y,dx*), *y* ∈ *R*^2^,*dx* ⊂ *R*^2^ for reconstruction which are supported over planes *Y* ⊂ *R*^2^ with uncertainty perpendicular to the line of flight but as well a second measurement the time-of-flight of the annihilating protons to the detectors [46]. Generally the point-spreads are modelled as cigar shaped two-dimensional Gaussians in the plane oriented by *N* angles *θ_n_* = 1,…, *N*, with high fidelity systems having *N* > 96, with standard deviation of uncertainty significantly larger along the lines of flight than perpendicular to them.

Classical parallel beam projection tomography reconstructs image planes *Y* ⊂ *R*^2^ via the Radon transform generated from sinograms indexed over a single space dimension *Z* ⊂ *R*^1^ arising from idealized line integrals [47]. Define the set of oriented lines in *R*^2^ parametetrized by their angles (*θ*) and offsets from the origin (*z*), *L_θ_*(*z*) = {(*y*^(1)^, *y*^(2)^)} ⊂ *R*^2^ with

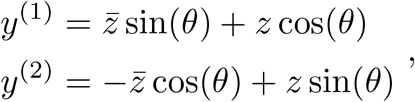

for indexing variable 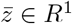 *R*^1^. To satisfy the basic sampling theorems for tomographic reconstruction [48], N sampling angles *θ_n_*, *n* = 1,…, *N*, akin to the sections in histology, are selected determined by the resolution required for the reconstruction. The point-spreads corresponding to the line integrals are 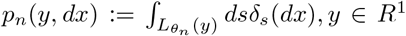, *dx* ⊂ *R*^2^, for *n* = 1,…, *N*. This gives the projections *P_n_I*(·) on R^1^ (see Supplementary Note S.1) with mean field of *J_n_*(·) the line integrals 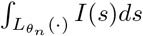 indexed over *R*^1^.

### 2.4 Integration of Tau Imaging Data into Multi-scale 3D Maps

We demonstrate, in this section, the efficacy of Projective LDDMM for aligning sets of 2D histology images to corresponding high field 3D MRI of MTL tissue taken postmortem from a brain sample with advanced AD (see Section 4.8 for details on specimen preparation). Additionally, we illustrate the benefits in using a measure-based framework to model microscopic data in this setting as it enables the estimation of distributions at varying scales and within particular volumes or 2D manifolds of interest.

We quantified relevant pathological measures in digital pathology images as counts of neurofibrillary tangles of hyperphosphorylated tau (NFTs) per cross-sectional area of tissue. NFTs were detected using a machine learningbased algorithm (see Section 4.5). Per pixel accuracy of identifying tau was evaluated with 10-fold cross validation, yielding an average AUC of 0.9860 and accuracy of 0.9729 (see Section 4.5 for individual fold metrics), and final counts of NFTs were validated against a reserved set of regions manually annotated for NFTs and totaling 25 million pixels.

Solutions to Variational Problem 2 (5d) yielded geometric reconstructions of histologically stained tissue in 3D. Figure 3 illustrates individual digitized sections on which NFTs were detected and the resulting positions of these slices following transformation via the geometric mappings to 3D were estimated. Coordination of both MRI and mapped histology with the Mai Paxinos Atlas [49] facilitated comparison of geometry between brain samples and is demonstrated in Figure 4, with coronal Mai views shown for an example intersecting histological slice.

**Fig. 3.**
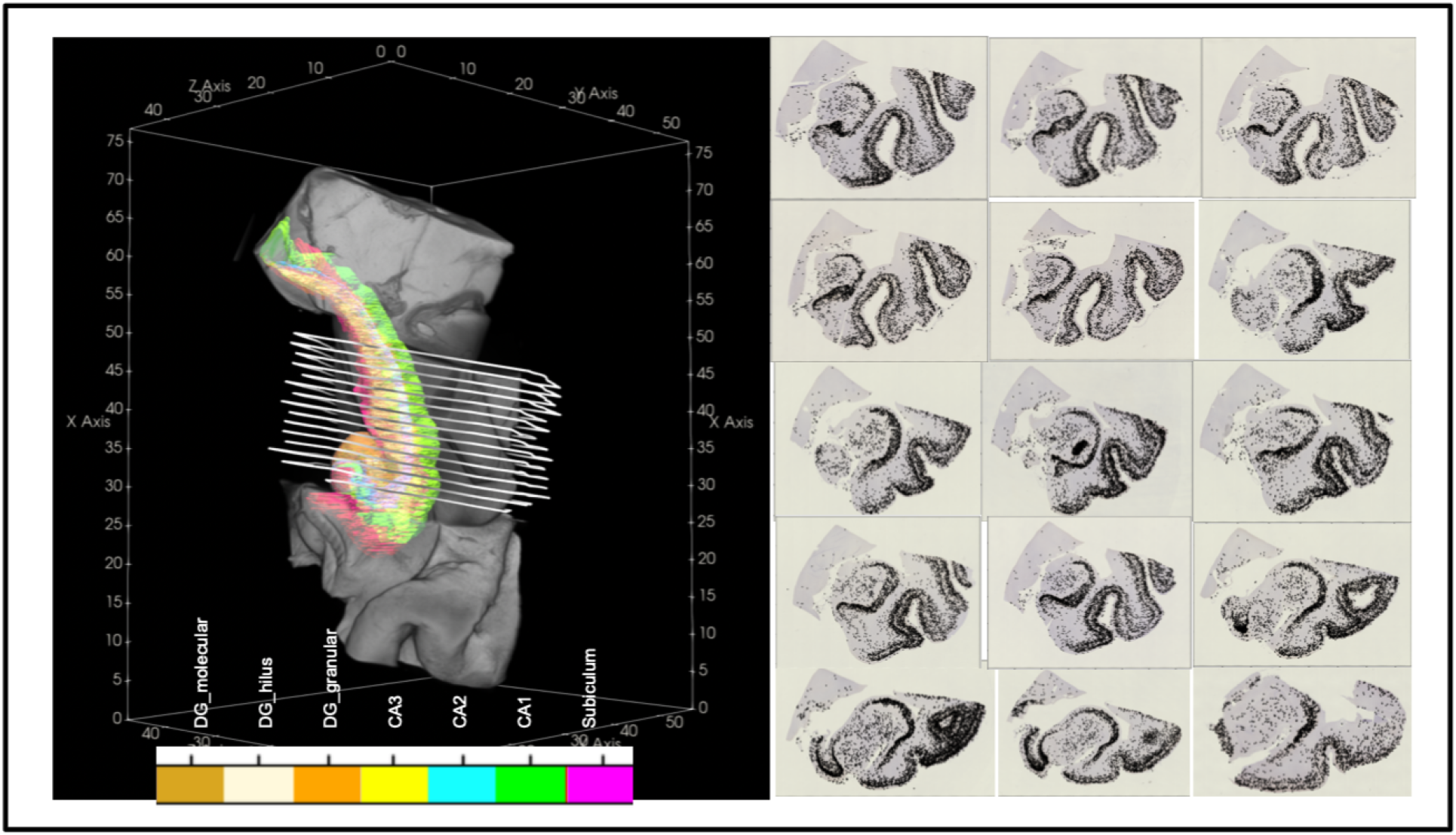
Complete set of PHF-1 stained histology sections for one of three blocks of MRI. 3D MRI shown with manual segmentations of MTL subregions (left). Boundary of each histological section on right outlined in white in position following transformation to 3D space (left). Detected tau tangles plotted as black dots over each histology slice (right).

**Fig. 4.**
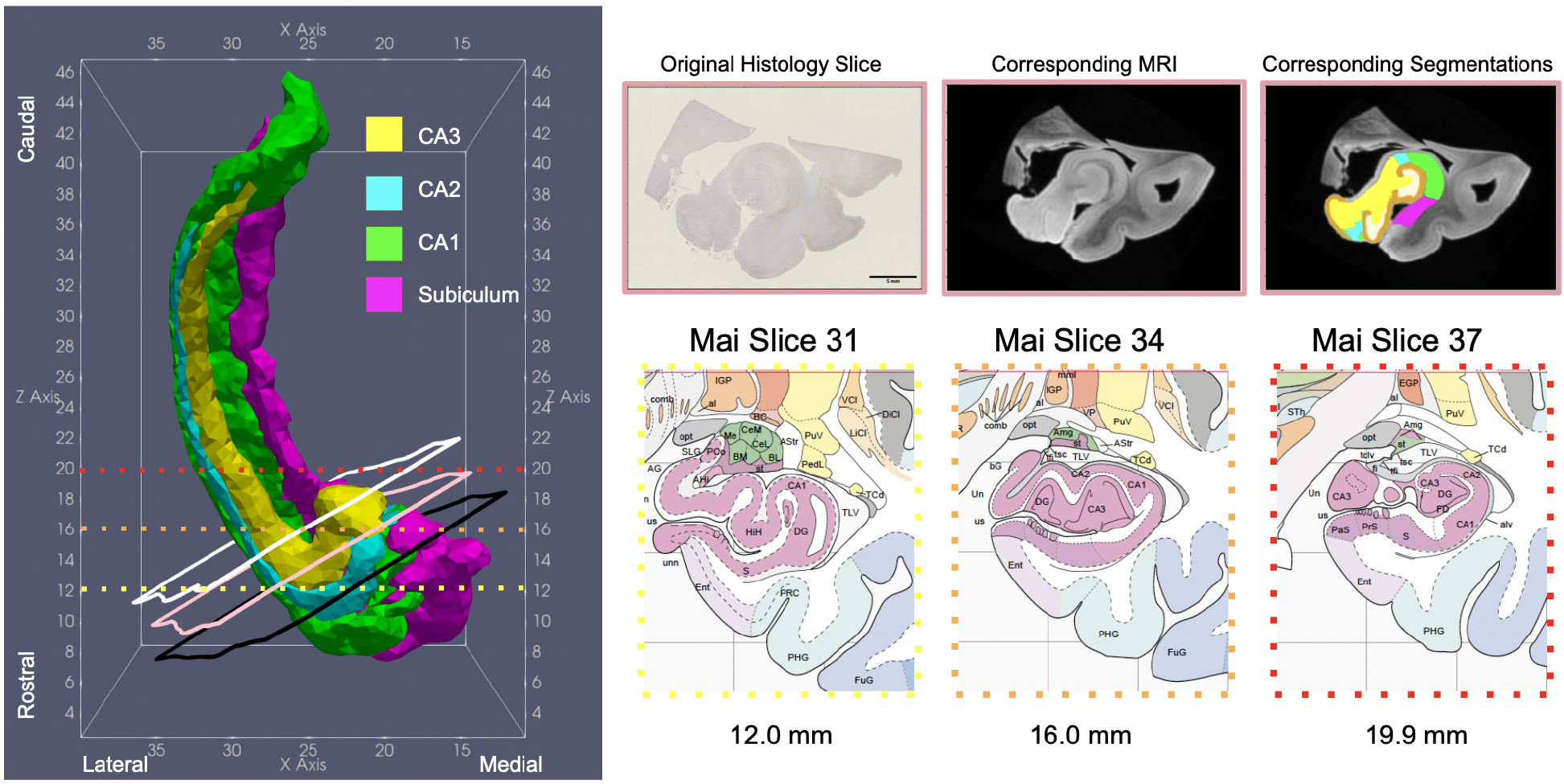
3D Reconstruction (left) of 4 MTL subregions for an advanced AD brain in the coordinate space of the Mai Paxinos Atlas. Corresponding section of histology and MRI (top right) shown and intersecting coronal planes taken from the pages of the Mai Atlas (bottom right).

Alignment accuracy was evaluated by comparison of manual segmentations on all histological images of one brain sample to those deformed from 3D MRI via estimated transformations. Figure 5 illustrates 4 representative comparisons. We quantified accuracy from these comparisons with Dice overlap and 95th percentile Hausdorff distance for MTL subregions of interest (see Supplementary Note S.2). The latter measure ranged from 1.2 mm to 1.8 mm across hippocampal subfields.

**Fig. 5.**
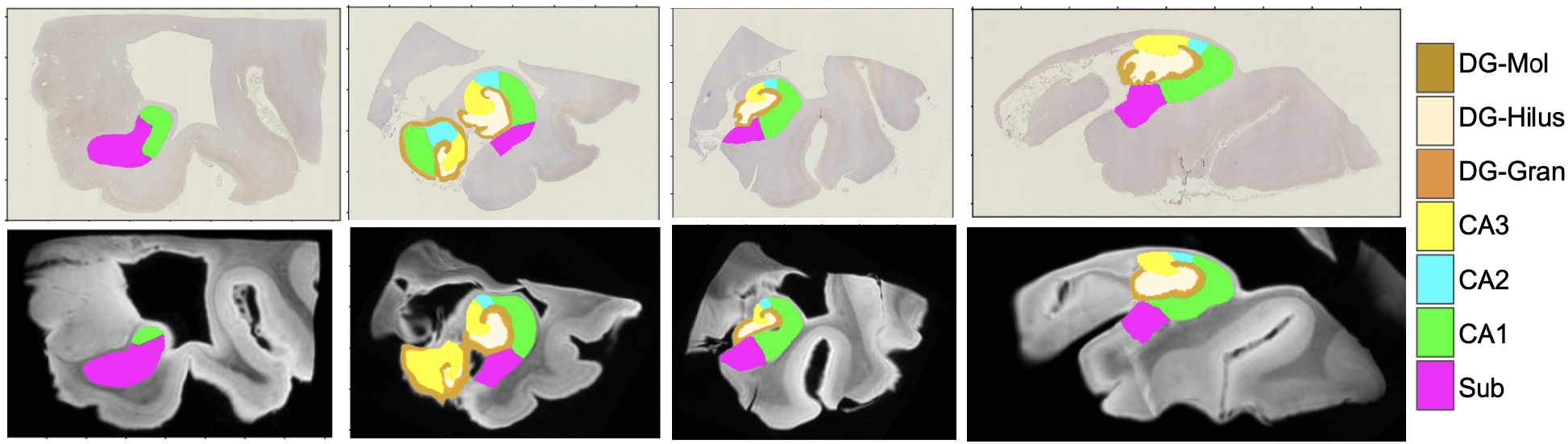
Selected histology slices with 2D segmentations (top row) ordered left to right as rostral to caudal. Corresponding MRI slices with 3D segmentations mapped to 2D via transformations *φ,ϕ_n_* (bottom row).

Counts of detected NFTs, cross-sectional tissue area, and MTL subregion (from MRI deformed to 2D) were computed in the space of histology slices. NFT densities (counts of NFTs per cross-sectional tissue area) were modeled as discrete particle measures and transported to the 3D space of the Mai Paxinos atlas via estimated transformations, *ϕ_n_, ϕ* (see Section 4.6). The flexibility afforded by a measure-based framework as in [36] is reflected in the diverse modes of resampling shown in Figure 6 both within volumes and over the surface of MTL subregions (see details in Supplementary Note S.4).

**Fig. 6.**
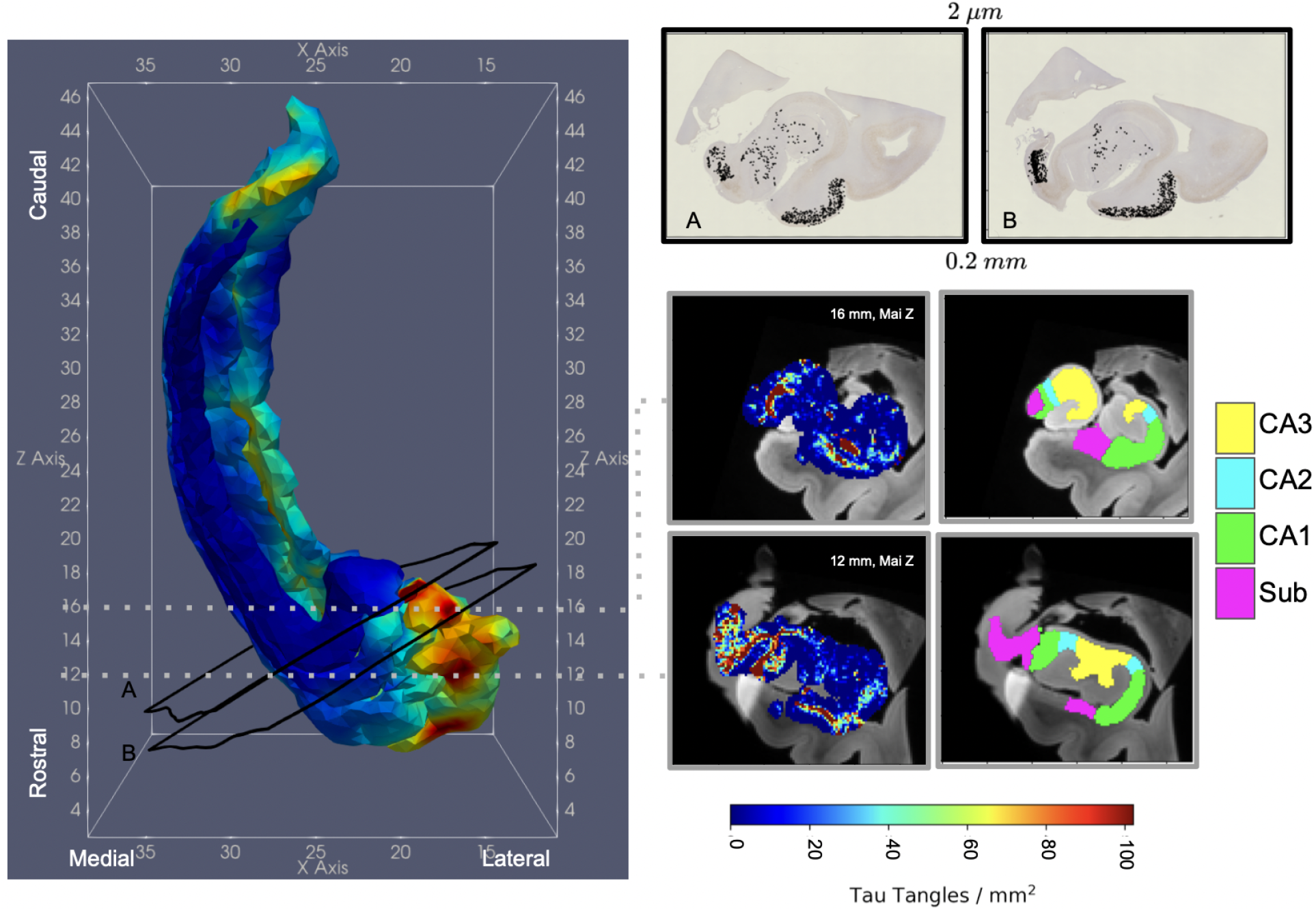
Distributions of NFT density within dense metric of hippocampus and over surface of hippocampal subregions (CA1, CA2, CA3, Subiculum).

Resampling via Gaussian kernels yields smoothed NFT densities computed within the dense metric (volume) of the brain at approximate resolutions of MRI. In contrast, spatial variations in NFT density within MTL subregions are visualized as smooth functions over the surface (2D manifold) of each corresponding region (see Section 4.7).

## 3 Discussion

The major contribution of this work is to extend the random orbit model of computational anatomy with added observation channels, representative of modern light sheet and optical imaging systems. In contrast to traditional 3D imaging modes (e.g. MRI), in these systems, observed measurements are not typically taken on the full domain of anatomy, putting them outside the traditional orbit of images generated by anatomical transformations. Here, we have focused on transferring digital pathology measures associated to neurodegenerative diseases to dense MR atlases, in which the observed histology are surjective measurements of the original 3D objects captured in MRI. We attempt to reconstruct the deformation of the template in 3D from the collection of 2D sections by way of a projection channel. The solving of this problem we call Projective LDDMM.

Simultaneously, we present a new way of expanding an orbit of images via the non-linear Scattering Transform of Mallat [28]. At the micro-scale, histology supports many multi-scale filtrations, each exhibiting textures of the tissue architecture. These filtrations form the basis of the expanded orbit imagery. We specifically describe a class of linear predictor models that generate a reduced dimension span from this basis to predict the MRI from the histology measurement. These predictors with the expanded orbit become the crucial link for estimating deformation of 3D MRI objects to match its conjectured 2D surjective measurements as manifest in the histology images. The power of the Scattering Transform is in its generation of filtrations across many scales, which are appropriate given the micro-scale histology which supports them.

Finally, we demonstrate a new class of Gaussian mixture models for modelling the random effects of tears and histological processing, each occurring independently in the histology coordinates per section. The mixtures are built to expand the orbit of images generated by the Scattering Transform by including non-foreground tissue as manifest by the smooth deformation of the MR template.

Appealing to the area of biomarker development in AD [31], we use the technologies of Projective LDDMM to reconstruct 3D maps of tau tangle density. Reconstruction within the dense 3D metric of the brain enables resampling of these measures at micron and millimeter resolution and within 3D volumes and 2D manifolds of interest. To visualize the tau densities in coordinates familiar to neuroscientists, we transfer the densities to the space of the Mai Paxinos atlas, which will enable comparison across brain samples in the future. We generate smoothed densities on the surface of hippocampal subregions by expanding tau measures in a respective basis for each surface. These bases are generated via the Laplace-Beltrami operator, moving from the classical Euclidean sines and cosines to complete ortho-normal bases for smooth curvilinear manifolds, such as these surfaces.

Similar to Yushkevich et. al [3], we found a strong spatial predominance of tau in the rostral third of the hippocampus we examined here. Together with variations in tau tangle density between hippocampal subregions, this predominance supports Braak’s initial observations [32, 33] and further underscores the spatial locality of AD. We are currently using our Projective LDDMM technologies to reconstruct tau profiles of an extended set of MTL regions including the entorhinal cortex and amygdala. By applying our methods to brain samples with intermediate and advanced AD, we plan to examine the specificity of such locality to AD and its temporal changes as compares to published trends in other imaging biomarkers.

## 4 Methods

### 4.1 Projective LDDMM Algorithm with In-Plane Transformation

To solve the Variational Problem 2 for a single 3D atlas, *I*_temp_, and a set of 2D targets (*J_*n*_*, for *n* = 1, …, *N*) we formulate an algorithm that alternately optimizes for the deformation in 3D space and the geometric transformation in 2D space, while holding the other fixed. The algorithm can be implemented to incorporate increasing complexity as needed first for crossing modalities and second for crossing resolutions, for instance, to map 3D MRI to 2D histological slices, as is presented here. In its simplest form, *I*_temp_ and *J* are of the same modality, yielding targets *J_n_* without expansion, and *ϕ_n_* modeled as a rigid motion in plane, as in Lee et. al [5], mapping histological sections to an atlas of the mouse brain. Transformations *φ, ϕ_n_* for *n* = 1, ⋯, *N* are estimated following Algorithm 1.

#### Algorithm 1

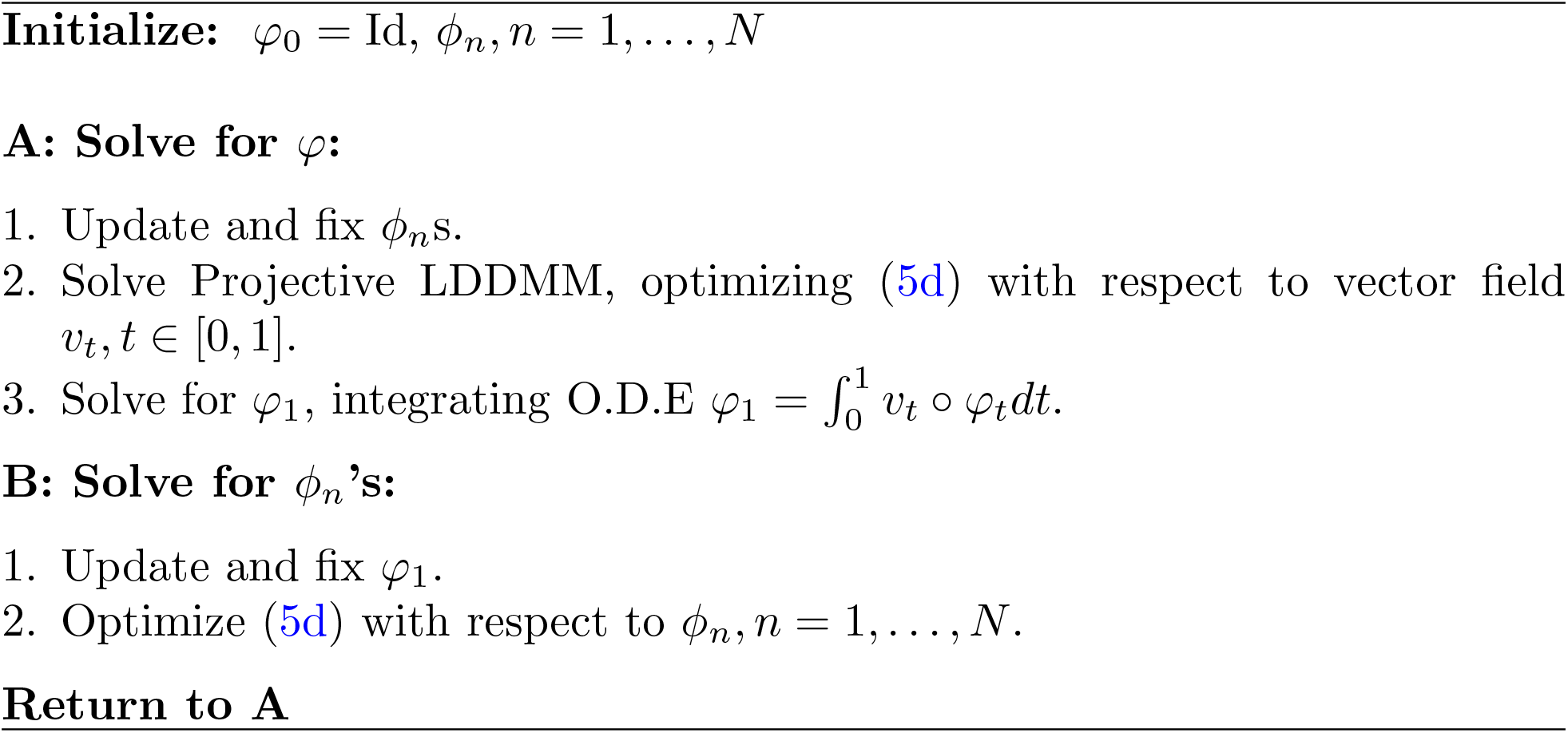

When *ϕ_n_*s are broadened to non-rigid diffeomorphisms, as in [6], each *ϕ_n_* is estimated in step B via a separate iteration of LDDMM for each target *J_n_*. Here, *ϕ_n_*s encapsulate both rigid and non-rigid components. Separate gradient based methods are used to update each component in step B with velocity fields updated using Hilbert gradient descent as in [50] and linear transform parameters updated by Gauss-Newton [51].

### 4.2 Linear Prediction Algorithm for Crossing Modalities with Scattering Transform

Crossing modalities at similar resolution (e.g. 3D MRI and downsampled 2D histology slices) requires a mapping between range spaces of template and target, giving a similar formulation to that used in Tward et. al [6]. In this work, we introduce the Scattering Transform [28] for crossing modalities at *differing* resolution whereby we build a predictive basis of images from our targets. We model contrast variations between histology and MRI by expanding the color space of our observed set of histology images {*J_n_*, *n* =1, ⋯, *N*} from three RGB dimensions to 48 feature dimensions reflecting local radiomic textures of histological scales.

Histology images are each resampled at the resolution of MRI to 48 “filtered images”, 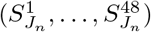 via Mallat’s Scattering Transform [28, 29] (see Supplementary Note S.3). We select a common 6-dimensional subspace of these 48 feature dimensions using PCA, yielding a basis of 6 discriminative “filtered images”, 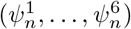 for each histology slice, *n* = 1,…, *N*, plus a constant image 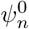. We predict the MRI contrast of our transformed projection as a linear combination of these basis elements 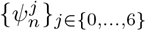 for each section *n* = 1, …, *N* with linear predictor:

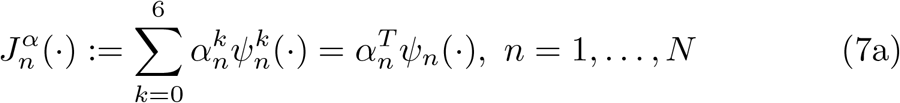

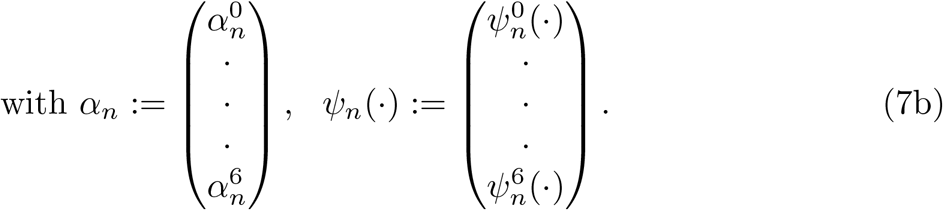

Figure 2 shows a mean field section using the scattering transform. These linear weights *α_n_* are estimated from initialized *ϕ_n_*s and *φ* following Algorithm 2. Initializations of *ϕ_n_* and *φ* are estimated following the approach in Tward et. al [27] in which cubic polynomials are used to match MRI range space to histology range space, generating solutions for *α_n_* via the pseudo-inverse:

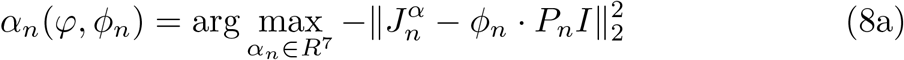

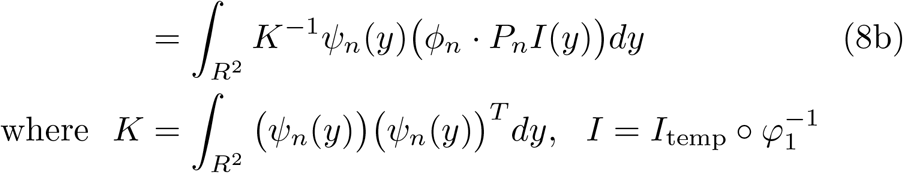

#### Algorithm 2

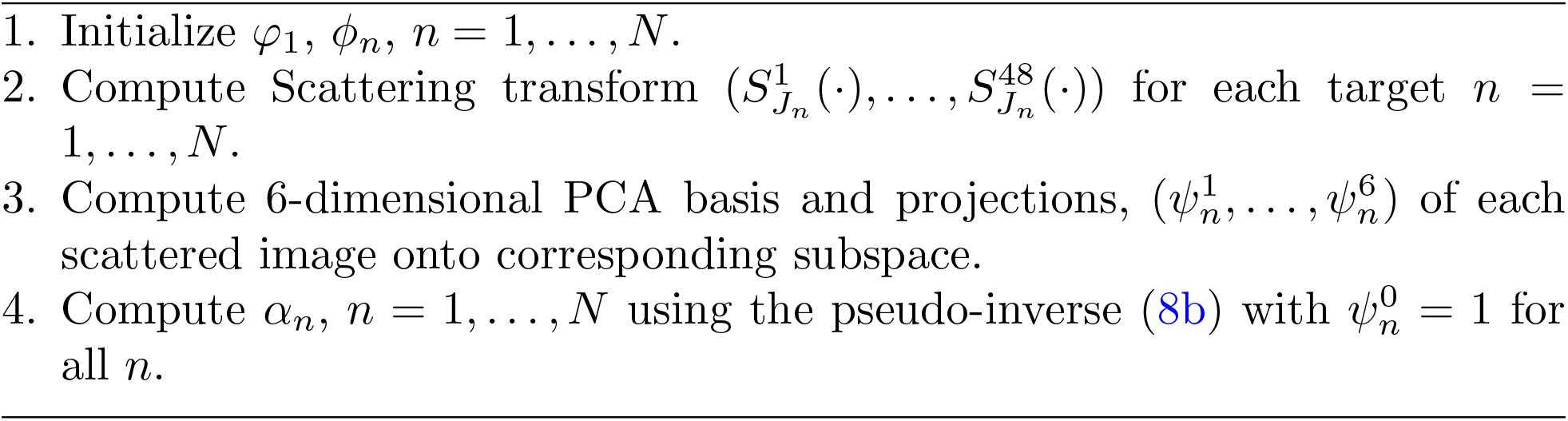

### 4.3 Projective LDDMM Algorithm with Crossing Modalities

The steps described in Algorithm 2 can be naturally incorporated into the framework of Projective LDDMM following Algorithm 3. Here, the linear prediction problem is solved using initial conditions for *ϕ_n_* and *φ*, and optimization of these geometric transformations then follows from a fixed set of estimated *α^n^*s.

#### Algorithm 3

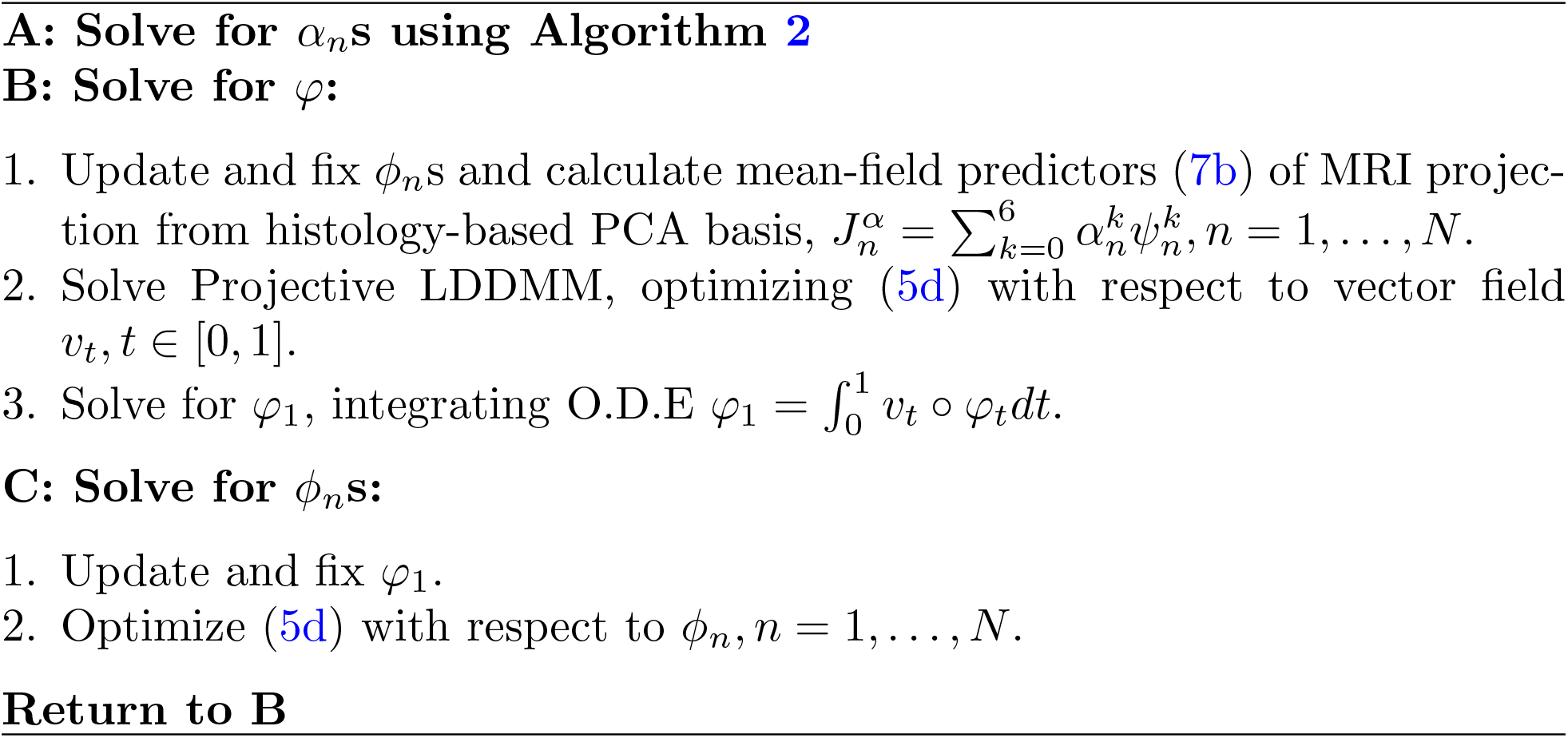

### 4.4 Projective LDDMM Algorithm with Multiple Models

Introduction of multiple models, as described in Section 2.2, replaces the matching cost in (5d) with that of (10a). As a result, the iteration in steps B-C of Algorithm 3 are replaced with an iterative algorithm based on the EM algorithm, implying it is monotonic in the cost. The complete-data likelihood for each histology plane *n* = 1, 2, … *N* as a function of parameters, *θ* = *ϕ_n_*, with 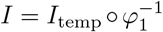 and *L*(·) mapping each location *y* ∈ *Y* to the set of labels {1, 2, 3} denoting foreground tissue, artifact, and background, is

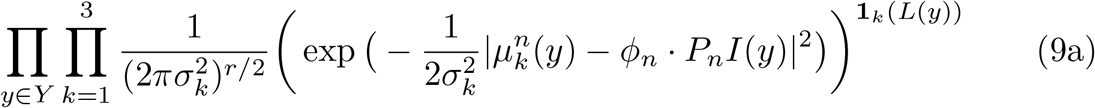

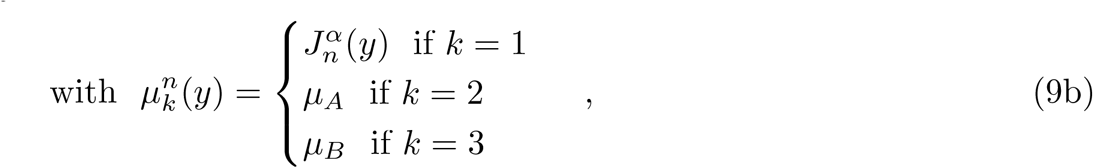

where *r* denotes the dimension of the range space of *I*(·) and *μ_A_* and *μ_B_* represent the means for artifact and background Gaussian distributions. The E-step takes the conditional expectation of the complete data log-likelihood with respect to the incomplete data, and the previous parameters *θ^old^*. The *M*-step generates our sequence of parameters:

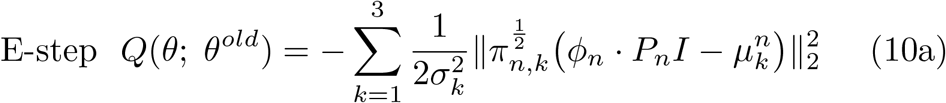

with 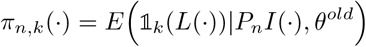.

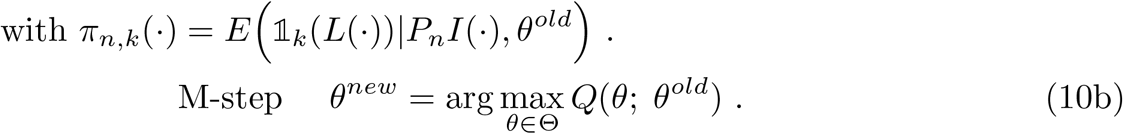

The spatial field of weights *π_n,k_* (·) is the conditional expectation of the indicator 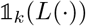. The Generalized EM (GEM) algorithm (see [30]) solves the maximization step: generating a sequence with increasing log-likelihood:

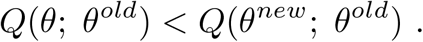

### 4.5 Tau Pathology Detection

Patterns of tau pathology are summarized as total counts of NFTs per mm^2^ of cross-sectioned tissue. NFT counts were computed using a 2-step algorithm: (1) prediction of per pixel probabilities of tau and (2) segmentation of these probability maps into discrete NFTs.

As described previously [52, 27], we used a convolutional neural network to model and predict probabilities of being part of a tau tangle for each pixel in a digital histology image. To capture larger contextual features as well as local information for producing per pixel probabilities at high resolutions, we trained UNETs [53] with the architecture described in Table 1. Training data was generated on every third slice of histology. Between 8 and 24 sample zones, sized 200-by-200 pixels were selected at random until 8 zones covered tissue (not background). Every pixel in each zone was manually annotated, 1 or 0, as part of a tau tangle or not. Estimates of accuracy in per pixel tau probabilities were computed using 10-fold cross validation on the entire training dataset. Table 2 shows accuracy metrics for each fold, with mean AUC and of 0.9860 and accuracy of 0.9729.

**Table 1.** Structure of UNET trained to detect tau tangles. Contraction layers are shown in the left 3 columns, and expansion layers in the right 3 columns. Number of parameters listed correspond to linear filters + bias vector. Conv: 3 × 3 convolution with stride 1, MP: 2 × 2 max pool, ReLU: Rectified Linear Unit, ConvT: 2 × 2 transposed convolution with stride 2. Note number of features doubles in the expansion layers due to concatenation with the contraction layers (skip connections).

| No. | Contract | Parameters | No. | Expand | Parameters |
| --- | --- | --- | --- | --- | --- |
| 1 | Conv | $8 \times 3 \times 3 \times 3 + 8$ | 18 | ConvT | $32 \times 64 \times 2 \times 2 + 32$ |
| 2 | Conv | $8 \times 8 \times 3 \times 3 + 8$ | 19 | Conv | $32 \times 64 \times 3 \times 3 + 32$ |
| 3 | MP | 0 | 20 | ReLU | 0 |
| 4 | Conv | $16 \times 8 \times 3 \times 3 + 16$ | 21 | Conv | $32 \times 32 \times 3 \times 3 + 32$ |
| 5 | ReLU | 0 | 22 | ReLU | 0 |
| 6 | Conv | $16 \times 16 \times 3 \times 3 + 16$ | 23 | ConvT | $16 \times 32 \times 2 \times 2 + 16$ |
| 7 | ReLU | 0 | 24 | Conv | $16 \times 32 \times 3 \times 3 + 16$ |
| 8 | MP | 0 | 25 | ReLU | 0 |
| 9 | Conv | $32 \times 16 \times 3 \times 3 + 32$ | 26 | Conv | $16 \times 16 \times 3 \times 3 + 16$ |
| 10 | ReLU | 0 | 27 | ReLU | 0 |
| 11 | Conv | $32 \times 32 \times 3 \times 3 + 32$ | 28 | ConvT | $8 \times 16 \times 2 \times 2 + 8$ |
| 12 | ReLU | 0 | 29 | Conv | $8 \times 16 \times 3 \times 3 + 8$ |
| 13 | MP | 0 | 30 | ReLU | 0 |
| 14 | Conv | $64 \times 32 \times 3 \times 3 + 64$ | 31 | Conv | $8 \times 8 \times 3 \times 3 + 8$ |
| 15 | ReLU | 0 | 32 | ReLU | 0 |
| 16 | Conv | $64 \times 64 \times 3 \times 3 + 64$ | 33 | Linear | $2 \times 8 + 2$ |
| 17 | ReLU | 0 |  |  |  |
|  |  |  |  | Total | 120,834 |

**Table 2.** 10-fold cross validation accuracy statistics for training data of brain sample.

| Trial | AUC | Precision | Recall | Accuracy |
| --- | --- | --- | --- | --- |
| 1 | 0.9997 | 0.0455 | 1.0000 | 0.9928 |
| 2 | 0.9983 | 0.0938 | 0.9917 | 0.9829 |
| 3 | 0.9963 | 0.1148 | 0.9846 | 0.9706 |
| 4 | 0.9984 | 0.2079 | 0.9971 | 0.9877 |
| 5 | 0.9796 | 0.3010 | 0.9242 | 0.9543 |
| 6 | 0.9915 | 0.0500 | 0.9620 | 0.9597 |
| 7 | 0.9865 | 0.2462 | 0.9031 | 0.9899 |
| 8 | 0.9239 | 0.0081 | 0.7079 | 0.9520 |
| 9 | 0.9989 | 0.1714 | 1.000 | 0.9939 |
| 10 | 0.9867 | 0.0255 | 0.9406 | 0.9454 |
| Average | 0.9860 | 0.1264 | 0.9411 | 0.9729 |

Counts of NFTs in each histology slice were generated by segmenting the probability maps output from the trained UNET. Segmentations were computed using an opensource implementation of the watershed algorithm [54] to extract connected components with “high probability” of tau. Each component was defined as an individual NFT, with center, area, and roundness computed as features.

### 4.6 Particle Representation of Histological Data

In the following two sections, we detail our means of representing histological data with a measure-based framework over physical space and feature space as introduced in [36]. We denote these measures, *μ*, borrowing the notation from [36] and describe the action of transformations on these measures to bring them to the 3D space of the Mai Paxinos Atlas and consequently our multiresolution resampling of them with a spatial kernel and feature map.

Here, we model histology data at the microscopic scale specifically following the discrete measure framework described in [36], where each particle of tissue, indexed by *i* ∈ *I*, carries a weighted Dirac measure over histology image space and a Dirac measure over the feature space *w_i_δ_y_i__* ⊗ *δ_f_i__*, *y_i_* ∈ *Y* ⊂ *R*^2^ and 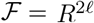. Weights reflect sampled tissue area captured in each particle measure. At the finest scale (*μ*^0^), this is defined as cross-sectional area in the histology plane *w_i_* ∈ {4*μm*^2^, 0}, computed with thresholding using Otsu’s method [55]. The first *ℓ* dimensions of 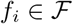 denote the number of tau tangles in the area of each of *ℓ* MTL subregions captured by the particle. The second *ℓ* dimensions denote the fraction of the particle’s total sampled tissue area (*w_i_*) within each of *ℓ* MTL subregions. At the finest scale, each particle captures one pixel of information, so the values of these features are 0 or 1, with

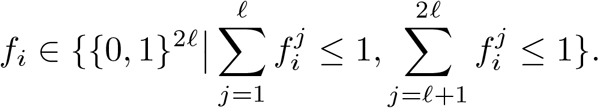

We transfer measures 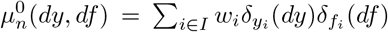 via diffeomorphisms *ϕ_n_* and *φ* and rigid transformation to the space of template *I* and the Mai Paxinos Atlas. Discrete weights *w_i_* adjust according to in plane expansion/contraction of cross-sectional tissue area, with adjustment at the fine scale by *ϕ_n_* given by the varifold action:

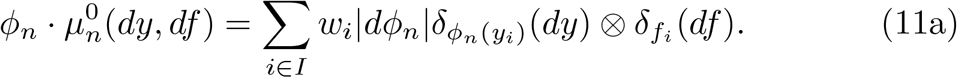

To cross scales we use the decomposition of the particle measures

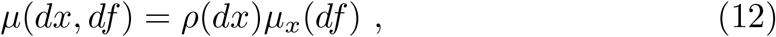

with *ρ* being the density of the model and *μ_x_* the field of conditional probabilities on the features. Our tranformation across scales non-linearly rescales space and smooths the empirical feature distributions on the features. Spatial resampling is determined by the function *π*(*x,x*′), which we define here to be the fraction the particle at *x* has assigned to it from the particle at *x*′, with *π*(*x,x*′)*dx*′ = 1. The smoothing on the field of conditional probabilities gives the remapped measure *μ*^1^.

Spatial resamplings at MRI resolutions (0.125 mm) and over surface boundaries of MTL subregions were achieved through isotropic Gaussian resampling and nearest neighbor resampling, respectively, through choice of *π* (see Supplementary Note S.4). Feature reduction occurs via maps 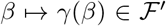 for probabilites *β*:

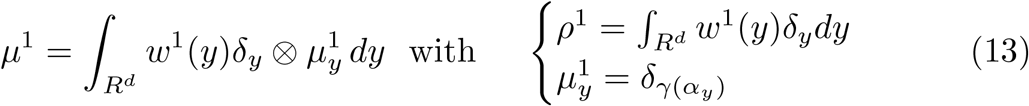

Here, *γ*(·) reduces feature dimension by taking empirical distributions *α_y_* over each of 2*ℓ* dimensions to expected first moments for each corresponding dimension, giving 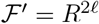 with:

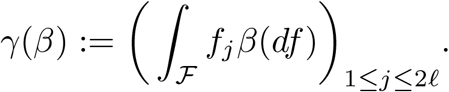

Total NFT density is computed from the sum of the first *ℓ* features while NFT density per region is computed from the ratio of feature value *j* to *ℓ* + *j* for any of *j* = 1, · · ·, **ℓ** MTL subregions.

### 4.7 Surface Smoothing with Laplace Beltrami Operator

Spatial variations in NFT density within MTL subregions are visualized as smooth functions over the surface of each corresponding region. Particle mass belonging to a given subregion volume is “projected” to the surface boundary using a nearest neighbor kernel for *π*(·, ·), as defined in Section 4.6:

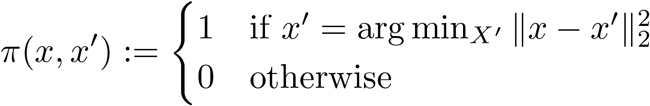

We construct functions, *g_τ_*(*x*) and *g_a_*(*x*), to represent the total number of NFTs and cross-sectional area of tissue from discrete particle measures of particles projected to the surface vertices *x_i_* ∈ *V*. To generate smooth representations of NFT density 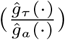, we build a complete orthonormal basis on each curved manifold using the Laplace-Beltrami operator [56], expanding each of the functions *g_τ_*(·) and *g_a_*(·) in this basis:

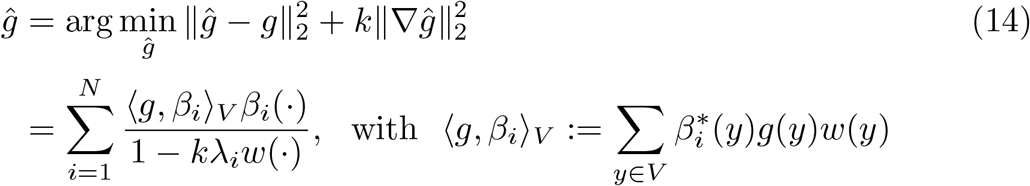

for both *g* = *g_τ_*(·) and *g* = *g_α_*(·), where 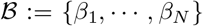 is a basis for the Laplace-Beltrami operator, and *k* the smoothing constant (see Supplementary Note S.5). We use these bases for smoothing over each corresponding surface in lieu of the standard Euclidean basis for *R*^3^, for which the Laplacian operator yields a basis of sines and cosines.

### 4.8 Specimen Preparation and Imaging

The brain tissue sample was prepared by the Johns Hopkins Brain Resource Center. From the formalin immersion fixed brain, a portion of the MTL including entorhinal cortex, amygdala, and hippocampus, was excised in 3 contiguous blocks of tissue, sized 20-30 mm in height and width, and 15 mm rostral-caudal (see reconstructed MRI of tissue blocks, Figure 3).

Each block was imaged with an 11T MR scanner at 0.125 mm isotropic resolution and then cut into two or three sets of 10 micron thick sections, spaced 1 mm apart. Each block yielded between 7 and 15 sections per set. Sets of sections were stained with PHF-1 for tau tangle detection, 6E10 for A*β* plaque detection, or Nissl, and digitized at 2 micron resolution.

### 4.9 Segmentations of MTL Subregions

MTL subregions within the hippocampus were manually delineated using Seg3D [57]. Individual block MRIs were rigidly aligned using an in-house manual alignment tool, and per voxel labels were saved for the composite MRI for each brain. Delineations were deduced from patterns of intensity differences, combined with previously published MR segmentations [58, 59, 60] and expert knowledge on the anatomy of the MTL. The established borders were applied in three other brains, showing consistent results (in preparation, EX, CC, SM, DT, JT, Alesha Seifert, Tilak Ratnanather, MA, MW, and MM). In the brain sample examined here, corresponding regional delineations were drawn on all histology sections stained with PHF-1 (see Figure 5). Delineations were based on visible anatomical markers and were afterwards confirmed with a corresponding Nissl-stained set of sections. In each of these sections, cytoarchitectonic borders between areas of the hippocampus were indicated, independently from the other datasets, using previously published cytoarchitectonic accounts of the MTL [61, 62, 63, 64, 65, 66]. Labels were assigned per pixel to 4x-downsampled histology images at a resolution of 32 microns and used to evaluate accuracy of registration (see Section 2.4). Particular regions of interest include cornu ammonis fields (CA1, CA2, CA3), and subiculum (see Figure 3).

## Supplementary information

This manuscript is accompanied by a set of Supplementary Notes (referenced S.1-S.8 in the main text).

## Acknowledgments

This research was funded in part by grants from the National Institutes of Health (U19-AG033655, P30-AG066507, P41-EB031771, R01-EB020062 (MM), T32-GM136577 (KS), U19-MH114821, R01-NS074980-10S1, RF1MH126732, RF1MH128875, RF1MH28888 (DT)), the Kavli Neuroscience Discovery Institute (MM,DT), and the Karen Toffler Charitable Trust (DT).

## Declarations

### Ethics Approval

This work was deemed not Human Subjects Research by the Johns Hopkins University IRB.

### Consent to Participate

Not applicable.

### Consent for Publication

Not applicable.

### Availability of Data and Materials

Due to size limitations, imaging data is available upon request.

### Code Availability

Code used for training and applying UNET in tau tangle detection can be found here: https://github.com/twardlab/ADproject. Code for solving Projective-LDDMM, building measure theoretic data representations, and resampling across scales can be found here: https://github.com/kstouff4/projective-lddmm.

### Authors’ Contributions

MW contrived the method for and oversaw manual segmentation of histology images and MRI. KS, MM, and DT designed and developed methods. KS and DT implemented algorithms. KS and MM drafted manuscript. All authors contributed to editing final manuscript.

## Supplementary Notes

### S.1 Classical Tomography

The Radon Transform is used in classical tomography [47] to describe the generation of sinograms as projected images at different angles. The transform is typically written in functional notation as an integral along one dimension, with Lebesgue measure as:

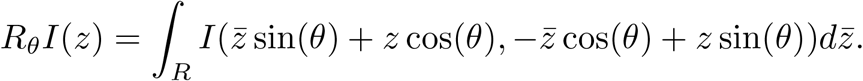

It can also be modeled as the integral over the line parametrized by two dimensions, an angle *θ* and affine offset *z* from the origin:

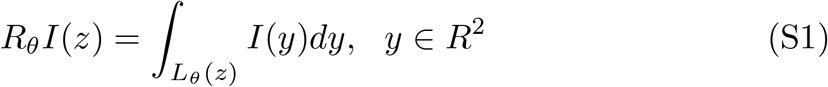

with the line defined by the paired angle and affine offset (*θ, z*) given explicitly by all *y* ∈ *R*^2^ such that:

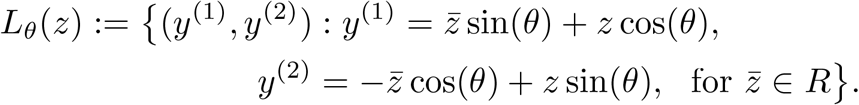

In the notation introduced in Section 2.1, we extend this to an integral over all of *R*^2^, using Dirac delta measures that assign nonzero measure only to lines in this same set:

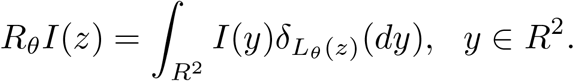

The line integral indexed by (*θ, z*) is modeled as a single projection *P_n_I*(*z*), *z* ∈ *R* indexed by a set of *n* = 1, …, *N* and is given as in (3b), with point spread, 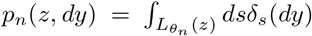, as defined in Section 2.3. We show below its equivalence to the line integral as defined in above (S1).

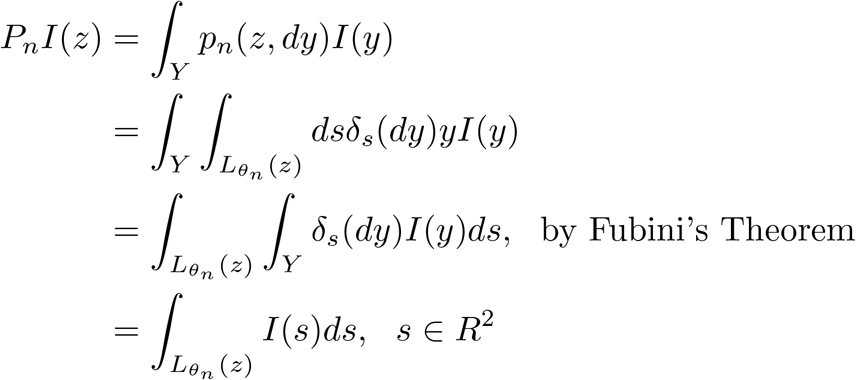

### S.2 Registration Accuracy

Similar to other groups [3], we evaluated accuracy of alignment between 2D histology and 3D MRI by looking at sets of discrete points (pixels) labeled in 2D versus corresponding voxels labeled in 3D and subsequently deformed to 2D (see Figure 5). Here, our sets of points were the sets of pixels (voxels) within a particular MTL subregion as delineated on 2D histology images and 3D MRI (see Section 4.9). We restricted our attention to particular subregions of interest (CA fields, dentate gyrus, and subiculum), and measured accuracy by Dice Score and 95th Percentile Hausdorff distance for each region on each slice of one brain. Average overlap scores were 0.65, 0.75, 0.72, 0.84 for subiculum, CA fields, dentate gyrus and whole hippocampus, respectively while average 95th percentile Hausdorff distance was 1.76 mm, 1.43 mm, 1.21 mm, and 1.69 mm for subiculum, CA fields, dentate gyrus, and whole hippocampus, respectively.

### S.3 Scattering Transform

The Scattering Transform, by Mallat and Bruna [28, 29], defines a cascade of alternating non-linear and non-commuting operators that generate from an image, *J_n_*(·) ∈ *L*^2^(*R^d^*), a set of “filtered images” 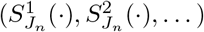 that capture textural information in the original image. Each “filtered image” is generated by a scattering propagator, *U* that takes the original image *J_n_* down a particular path, *p*, of alternating convolutions with wavelets (localized waveforms) and modulus operations. The path, *p*, is defined by a set of parameters 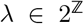 that scale a mother wavelet, *ζ*, so as to capture lower and lower frequency information.

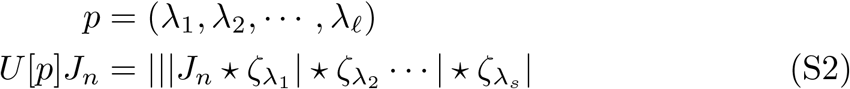

In our setting, we compute a subsampled Scattering Transform, 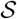, of each of our histology images, using an algorithm similar to the “Filterbank” algorithm [67, 68] in which images are downsampled in parallel with scattering. The path dependent propagator corresponding to 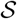 is *U_s_*[·]:

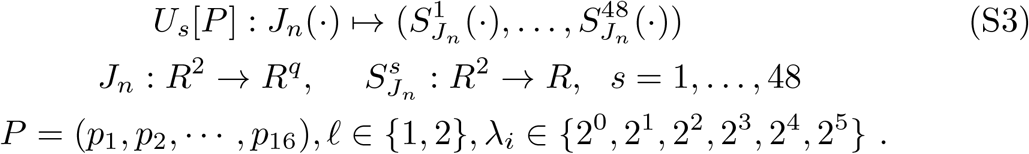

We use 16 paths, *p_i_* ∈ *P* of length 1 or 2 and a high pass Gaussian filter, with width dilated according to *λ_i_*, in place of a traditional wavelet to achieve a representation both translation and rotation invariant in addition to Lipschitz continuous to small deformations. Each of the R,G,B channels of histology images are propagated independently along the same paths. Histology images are downsampled by a factor of 32 to reach the approximate resolution of MRI. Together, the subsampling and scattering of each channel yield a total of 48 scattering coefficients for each pixel in the downsampled histology image, or 48 “filtered images” per target histology image.

### S.4 Resampling in Mai Atlas Space

To compute and compare NFT density measures across brain samples, we rigidly aligned all samples to the reference brain in the Mai Paxinos Atlas [49]. We used a manual alignment tool, created in-house, to select optimal alignments between surface renderings of the Hippocampus, Amygdala, and ERC of our brain samples and that of the Mai brain.

Distributions of NFT density were computed in the coordinates of the Mai atlas, according to choice of *π*(*x, x*′) governing physical spatial spread, and *γ*(·) governing smoothing over conditional feature distributions (13). In all cases, total mass (2D cross-sectional tissue area of histological images) and total number of NFTs were conserved. To achieve this, initial NFT feature values (counts per MTL subregion) were reformulated following physical transformation (11a) as counts of NFTs per MTL subregion per weight of particle (i.e. total cross-sectional area):

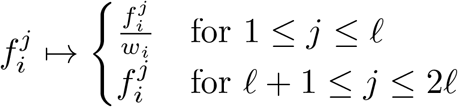

We highlight three different modes of resampling. Volumetric resampling (e.g. at mm resolution) was computed with a 3D isotropic Gaussian kernel with width, *σ*, and with new particles in a regular lattice, *x*′ ∈ *X*′.

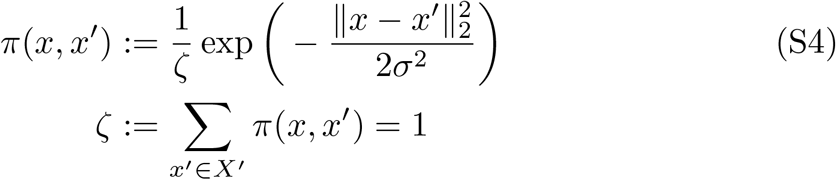

Resampling over 2D manifolds (e.g. the surface of ERC, Amygdala, CA1, or Subiculum) was computed using a nearest neighbor kernel, assigning all weight (tissue area) and NFTs from a particle at the fine scale to a single particle on the 2D manifold (e.g. vertex of a triangular mesh).

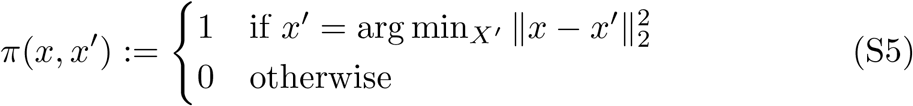

Finally, resampling to a regular 1D lattice (e.g. the rostral-caudal axis of the human brain) was computed using an anisotropic Gaussian kernel to spread particle mass widely in two dimensions and narrowly in the third, with dimensions treated independently.

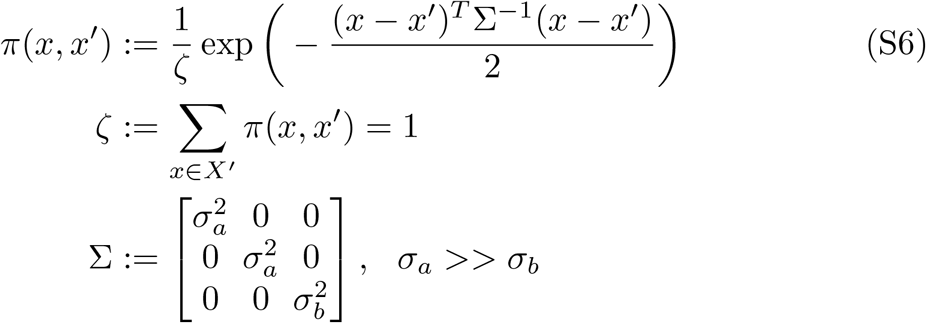

In each case, feature reduction occurred via computation of expected first moments, as described in section 4.6.

### S.5 Laplace Beltrami Smoothing Solution

The variational solution to (14) is given by:

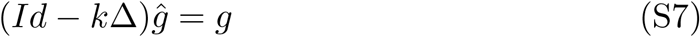

where Δ is a Laplacian operator. Here, we take Δ as the Laplace Beltrami operator and compute an eigenbasis 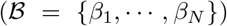 and eigenvalues ({λ_1_, ⋯, λ_*N*_}) via the Finite Elements Method (FEM) [56]. Expansion of (S7) in this eigenbasis yields smoothed 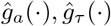 for choice of parameter k:

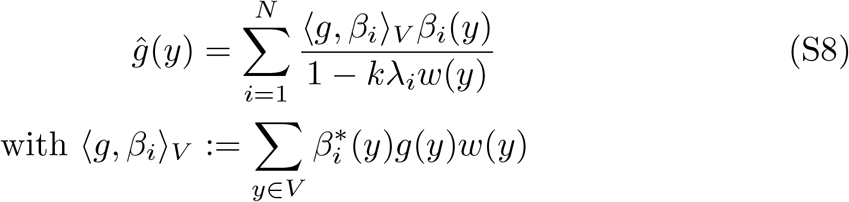

Both 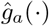 and 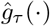 are normalized independently so total cross sectional area and numbers of NFTs projected to the surface are conserved before and after smoothing. NFT densities are computed as the ratio of the normalized, smoothed functions: 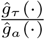 and plotted over the surfaces of given MTL subregions (see Figure 6).

## References

[1] Gupta, R., Kurc, T., Saltz, J.H.: Introduction to Digital Pathology from Historical Perspectives to Emerging Pathomics. https://doi.org/10.1007/978-3-030-83332-9-1

[2] Lu, C., Shiradkar, R., Liu, Z.: Integrating pathomics with radiomics and genomics for cancer prognosis: A brief review. Chinese Journal of Cancer Research 33, 563–573 (2021). https://doi.org/10.21147/j.issn.1000-9604.2021.05.03

[3] Yushkevich, P.A., de Onzoño Martin, M.M.I., Ittyerah, R., Lim, S., Lavery, M., Wang, J., Hung, L.Y., Vergnet, N., Ravikumar, S., Xie, L., Dong, M., DeFlores, R., Cui, S., McCollum, L., Ohm, D.T., Robinson, J.L., Schuck, T., Grossman, M., Tisdall, M.D., Prabhakaran, K., Mizsei, G., Das, S.R., Artacho-Pérula, E., del Mar Arroyo Jiménez, M., López, M.M., Rabal, M.P.M., Romero, F.J.M., Lee, E.B., Trojanowski, J.Q., Wisse, L.E.M., Wolk, D.A., Irwin, D.J., Insausti, R.: 3d mapping of tau neurofibrillary tangle pathology in the human medial temporal lobe. In: 2020 IEEE 17th International Symposium on Biomedical Imaging (ISBI), pp. 1312–1316 (2020). https://doi.org/10.1109/ISBI45749.2020.9098462

[4] Goubran, M., Leuze, C., Hsueh, B., Aswendt, M., Ye, L., Tian, Q., Cheng, M.Y., Crow, A., Steinberg, G.K., McNab, J.A., Deisseroth, K., Zeineh, M.: Multimodal image registration and connectivity analysis for integration of connectomic data from microscopy to mri. Nature Communications 10 (2019). https://doi.org/10.1038/s41467-019-13374-0

[5] Lee, B.C., Tward, D.J., Mitra, P.P., Miller, M.I.: On variational solutions for whole brain serial-section histology using a Sobolev prior in the computational anatomy random orbit model. PLoS Computational Biology 14(12), 1–20 (2018). https://doi.org/10.1371/journal.pcbi.1006610

[6] Tward, D., Li, X., Huo, B., Lee, B., Mitra, P., Miller, M.: 3D Mapping of Serial Histology Sections with Anomalies Using a Novel Robust Deformable Registration Algorithm 3D Mapping of Serial Sections via Robust Deformable Registration 163. In: LNCS, vol. 11846, pp. 162–173 (2019). https://doi.org/10.1007/978-3-030-33226_18

[7] Lee, B.C., Lin, M.K., Fu, Y., Hata, J., Miller, M.I., Mitra, P.P.: Multimodal cross-registration and quantification of metric distortions in marmoset whole brain histology using diffeomorphic mappings. Journal of Comparative Neurology 529(2), 281–295 (2021) https://arxiv.org/abs/arXiv:1805.04975v2. https://doi.org/10.1002/cne.24946

[8] Iglesias, J.E., Insausti, R., Lerma-Usabiaga, G., Bocchetta, M., Van Leemput, K., Greve, D.N., van der Kouwe, A., Fischl, B., Caballero-Gaudes, C., Paz-Alonso, P.M.: A probabilistic atlas of the human thalamic nuclei combining ex vivo MRI and histology. NeuroImage 183(July), 314–326 (2018) https://arxiv.org/abs/1806.08634. https://doi.org/10.1016/j.neuroimage.2018.08.012

[9] Perens, J., Salinas, C.G., Skytte, J.L., Roostalu, U., Dahl, A.B., Dyrby, T.B., Wichern, F., Barkholt, P., Vrang, N., Jelsing, J., Hecksher-Sørensen, J.: An Optimized Mouse Brain Atlas for Automated Mapping and Quantification of Neuronal Activity Using iDISCO+ and Light Sheet Fluorescence Microscopy. Neuroinformatics 19(3), 433–446 (2021). https://doi.org/10.1007/s12021-020-09490-8

[10] Hillman, E.M.C., Voleti, V., Li, W., Yu, H.: Light-Sheet Microscopy in Neuroscience. Annual Review of Neuroscience 42, 295–313 (2019). https://doi.org/10.1146/annurev-neuro-070918-050357

[11] Veldman, M.B., Park, C.S., Eyermann, C.M., Zhang, J.Y., Zuniga-Sanchez, E., Hirano, A.A., Daigle, T.L., Foster, N.N., Zhu, M., Langfelder, P., Lopez, I.A., Brecha, N.C., Zipursky, S.L., Zeng, H., Dong, H.W., Yang, X.W.: Brainwide genetic sparse cell labeling to illuminate the morphology of neurons and glia with cre-dependent morf mice. Neuron 108, 111–1276 (2020). https://doi.org/10.1016/j.neuron.2020.07.019

[12] Bergenstråhle, J., Larsson, L., Lundeberg, J.: Seamless integration of image and molecular analysis for spatial transcriptomics workflows. BMC Genomics 21(1), 1–7 (2020). https://doi.org/10.1186/s12864-020-06832-3

[13] Chen, W.T., Lu, A., Craessaerts, K., Pavie, B., Sala Frigerio, C., Corthout, N., Qian, X., Laláková, J., Kühnemund, M., Voytyuk, I., Wolfs, L., Mancuso, R., Salta, E., Balusu, S., Snellinx, A., Munck, S., Jurek, A., Fernandez Navarro, J., Saido, T.C., Huitinga, I., Lundeberg, J., Fiers, M., De Strooper, B.: Spatial Transcriptomics and In Situ Sequencing to Study Alzheimer’s Disease. Cell 182(4), 976–99119 (2020). https://doi.org/10.1016/j.cell.2020.06.038

[14] Miller, M.I., Fan, J., Tward, D.J.: Multi scale diffeomorphic metric mapping of spatial transcriptomics datasets. In: 2021 IEEE/CVF Conference on Computer Vision and Pattern Recognition Workshops (CVPRW), pp. 4467–4475 (2021). https://doi.org/10.1109/CVPRW53098.2021.00504

[15] Palla, G., Fischer, D.S., Regev, A., Theis, F.J.: Spatial components of molecular tissue biology. Nature Biotechnology (2022). https://doi.org/10.1038/s41587-021-01182-1

[16] Grenander, U., Miller, M.I.: Computational Anatomy: An Emerging Discipline. Applied Mathematics 56(4), 617–694 (1998)

[17] Grenander, U., Miller, M.I.: Pattern Theory: From Representation To Inference, pp. 1–596. Oxford University Press, New York (2007)

[18] Miller, M.I., Younes, L., Trouvé, A.: Diffeomorphometry and geodesic positioning systems for human anatomy. Technology 02(01), 36–43 (2014). https://doi.org/10.1142/s2339547814500010

[19] Miller, M.I., Arguillère, S., Tward, D.J., Younes, L.: Computational anatomy and diffeomorphometry: A dynamical systems model of neuroanatomy in the soft condensed matter continuum. Wiley Interdisciplinary Reviews: Systems Biology and Medicine 10(6), 1–42 (2018). https://doi.org/10.1002/wsbm.1425

[20] Miller, M.I., Younes, L.: Group actions, homeomorphisms, and matching: A general framework. International Journal of Computer Vision 41, 61–84 (2001)

[21] Miller, M.I., Trouvá, A., Younes, L.: On the metrics and Euler-Lagrange equations of computational anatomy. Annual Review of Biomedical Engineering 4, 375–405 (2002). https://doi.org/10.1146/annurev.bioeng.4.092101.125733

[22] Avants, B.B., Epstein, C.L., Grossman, M., Gee, J.C.: Symmetric diffeomorphic image registration with cross-correlation: Evaluating automated labeling of elderly and neurodegenerative brain. Medical Image Analysis 12(1), 26–41 (2008). https://doi.org/10.1016/j.media.2007.06.004

[23] Christensen, G.E., Johnson, H.J.: Consistent image registration. IEEE Transactions on Medical Imaging 20(7), 568–582 (2001). https://doi.org/10.1109/42.932742

[24] Avants, B.B., Epstein, C.L., Grossman, M., Gee, J.C.: Symmetric Diffeomorphic Image Registration with Cross-Correlation: Evaluating Automated Labeling of Elderly and Neurodegenerative Brain

[25] Pluim, J.P.W., Maintz, J.B.A.A., Viergever, M.A.: Mutual-information-based registration of medical images: A survey. IEEE Transactions on Medical Imaging 22(8), 986–1004 (2003). https://doi.org/10.1109/TMI.2003.815867

[26] Heinrich, M.P., Jenkinson, M., Bhushan, M., Matin, T., Gleeson, F.V., Brady, S.M., Schnabel, J.A.: MIND: Modality independent neighbourhood descriptor for multi-modal deformable registration. Medical Image Analysis 16(7), 1423–1435 (2012). https://doi.org/10.1016/j.media.2012.05.008

[27] Tward, D., Brown, T., Kageyama, Y., Patel, J., Hou, Z., Mori, S., Albert, M., Troncoso, J., Miller, M.: Diffeomorphic Registration With Intensity Transformation and Missing Data: Application to 3D Digital Pathology of Alzheimer’s Disease. Front. Neurosci. 14(February), 1–18 (2020). https://doi.org/10.3389/fnins.2020.00052

[28] Mallat, S.: Group invariant scattering. Commun. Pur. Appl. Math. 65(10), 1331–1398 (2012)

[29] Bruna, J., Mallat, S.: Invariant scattering convolution networks. IEEE Trans. Pattern Anal. Mach. Intell. 35(8), 1872–1886 (2013) https://arxiv.org/abs/1203.1513. https://doi.org/10.1109/TPAMI.2012.230

[30] Dempster, A.P., Laird, N.M., Rubin, D.B.: Maximum likelihood from incomplete data via the em algorithm. J. R. Stat. Soc. Series B 39(1), 1–38 (1977)

[31] Jack, C.R., Bennett, D.A., Blennow, K., Carrillo, M.C., Dunn, B., Haeberlein, S.B., Holtzman, D.M., Jagust, W., Jessen, F., Karlawish, J., Liu, E., Molinuevo, J.L., Montine, T., Phelps, C., Rankin, K.P., Rowe, C.C., Scheltens, P., Siemers, E., Snyder, H.M., Sperling, R., Elliott, C., Masliah, E., Ryan, L., Silverberg, N.: NIA-AA Research Framework: Toward a biological definition of Alzheimer’s disease. Alzheimer’s Dement. 14(4), 535–562 (2018). https://doi.org/10.1016/j.jalz.2018.02.018

[32] Braak, H., Braak, E.: Neuropathological stageing of Alzheimer-related changes. Acta Neuropathol. 82(4), 239–259 (1991). https://doi.org/10.1007/BF00308809

[33] Braak, H., Alafuzoff, I., Arzberger, T., Kretzschmar, H., Tredici, K.: Staging of Alzheimer disease-associated neurofibrillary pathology using paraffin sections and immunocytochemistry. Acta Neuropathologica 112(4), 389–404 (2006). https://doi.org/10.1007/s00401-006-0127-z

[34] Hyman, B.T., Phelps, C.H., Beach, T.G., Bigio, E.H., Cairns, N.J., Carrillo, M.C., Dickson, D.W., Duyckaerts, C., Frosch, M.P., Masliah, E., Mirra, S.S., Nelson, P.T., Schneider, J.A., Thal, D.R., Thies, B., Trojanowski, J.Q., Vinters, H.V., Montine, T.J.: National Institute on Aging-Alzheimer’s Association guidelines for the neuropathologic assessment of Alzheimer’s disease. Alzheimer’s and Dementia 8(1), 1–13 (2012). https://doi.org/10.1016/J.JALZ.2011.10.007

[35] Kulason, S., Tward, D.J., Brown, T., Sicat, C.S., Liu, C.F., Ratnanather, J.T., Younes, L., Bakker, A., Gallagher, M., Albert, M., Miller, M.I.: Cortical thickness atrophy in the transentorhinal cortex in mild cognitive impairment. NeuroImage: Clin. 21, 101617 (2019). https://doi.org/10.1016/j.nicl.2018.101617

[36] Miller, M.I., Tward, D., Trouvé, A.: Molecular Computational Anatomy: A Unified Molecular to Tissue Continuum Via Measure Representations. BME Frontiers (in press)

[37] Tournier, J.-D., Mori, S., Leemans, A., Morgan, R.H., Reson, M., Author, M.: Diffusion Tensor Imaging and Beyond NIH Public Access Author Manuscript. Magn Reson Med 65(6), 1532–1556 (2011). https://doi.org/10.1002/mrm.22924.Diffusion

[38] Bracewell, R.N.: Strip Integration in Radio Astronomy. Australian Journal of Physics 9, 198 (1956). https://doi.org/10.1071/PH560198

[39] Zhao, S.-R., Halling, H.: A new fourier method for fan beam reconstruction. In: 1995 IEEE Nuclear Science Symposium and Medical Imaging Conference Record, vol. 2, pp. 1287–12912 (1995). https://doi.org/10.1109/NSSMIC.1995.510494

[40] Dupuis, P., Grenander, U., Miller, M.I.: Variational problems on flows of diffeomorphisms for image matching. Quarterly of Applied Mathematics 56(3), 587–600 (1998)

[41] Tang, X., Oishi, K., Faria, A.V., Hillis, A.E., Albert, M.S., Mori, S., Miller, M.I.: Bayesian parameter estimation and segmentation in the multi-atlas random orbit model. PLoS ONE 8 (2013). https://doi.org/10.1371/journal.pone.0065591

[42] Wu, D., Ma, T., Ceritoglu, C., Li, Y., Chotiyanonta, J., Hou, Z., Hsu, J., Xu, X., Brown, T., Miller, M.I., Mori, S.: Resource atlases for multi-atlas brain segmentations with multiple ontology levels based on t1-weighted mri. NeuroImage 125, 120–130 (2016). https://doi.org/10.1016/j.neuroimage.2015.10.042

[43] Doshi, J., Erus, G., Ou, Y., Resnick, S.M., Gur, R.C., Gur, R.E., Satterthwaite, T.D., Furth, S., Davatzikos, C.: Muse: Multi-atlas region segmentation utilizing ensembles of registration algorithms and parameters, and locally optimal atlas selection. NeuroImage 127, 186–195 (2016). https://doi.org/10.1016/j.neuroimage.2015.11.073

[44] Joshi, S., Miller, M.I.: Maximum a posteriori estimation with Good’s roughness for three-dimensional optical-sectioning microscopy. J. Opt. Soc. Am. A 10(5), 1078–1085 (1993). https://doi.org/10.1364/JOSAA.10.001078

[45] Gibson, S.F., Lanni, F.: Diffraction by a circular aperture as a model for three-dimensional optical microscopy. J. Opt. Soc. Am. A 6(9), 1357–1367 (1989). https://doi.org/10.1364/JOSAA.6.001357

[46] Snyder, D., Thomas Jr., L., Ter-Pogossian, M.: Mathematical model for positron-emission tomography systems having time-of-flight measurements. IEEE Transactions on Nuclear Science NS-28(3), 3575–83 (1981)

[47] Barrett, H.H.: Iii the radon transform and its applications. Progress in Optics, vol. 21, pp. 217–286. Elsevier (1984). https://doi.org/10.1016/S0079-6638(08)70123-9. https://www.sciencedirect.com/science/article/pii/S0079663808701239

[48] Snyder, D., Cox, J.: An overview of reconstruction tomography and limitations imposed by a finite number of projections. In: Proceedings of Workshop on Reconstruction Tomography in Diagnostic Radiology and Nuclear Medicine, Puerto Rico (1975)

[49] Mai, J.K., Paxinos, G., Voss, T.: Atlas of the Human Brain, 3rd edn. Elsevier Inc, New York (2008)

[50] Beg, M.F., Miller, M.I., Trouvé, A., Younes, L.: Computing large deformation metric mappings via geodesic flows of diffeomorphisms. Int. J. Comput. Vis. 61(2), 139–157 (2005). https://doi.org/10.1023/B:VISI.0000043755.93987.aa

[51] Tward, D.J.: An optical flow based left-invariant metric for natural gradient descent in affine image registration. Frontiers in Applied Mathematics and Statistics, 61

[52] Stouffer, K.M., Wang, Z., Xu, E., Lee, K., Lee, P., Miller, M.I., Tward, D.J.: From picoscale pathology to decascale disease: Image registration with a scattering transform and varifolds for manipulating multiscale data. In: Syeda-Mahmood, T., Li, X., Madabhushi, A., Greenspan, H., Li, Q., Leahy, R., Dong, B., Wang, H. (eds.) Multimodal Learning for Clinical Decision Support, pp. 1–11. Springer, Cham (2021)

[53] Ronneberger, O., Fischer, P., Brox, T.: U-net: Convolutional networks for biomedical image segmentation. CoRR abs/1505.04597 (2015) https://arxiv.org/abs/1505.04597

[54] Bradski, G.: The OpenCV Library. Dr. Dobb’s J. Softw. Tools (2000)

[55] Otsu, N.: A threshold selection method from gray-level histograms. IEEE Transactions on Systems, Man, and Cybernetics 9(1), 62–66 (1979). https://doi.org/10.1109/TSMC.1979.4310076

[56] Qiu, A., Bitouk, D., Miller, M.I.: Smooth functional and structural maps on the neocortex via orthonormal bases of the laplace-beltrami operator. IEEE Transactions on Medical Imaging 25(10), 1296–1306 (2006). https://doi.org/10.1109/TMI.2006.882143

[57] CIBC Seg3D: Volumetric Image Segmentation and Visualization. Scientific Computing and Imaging Institute (SCI), Download from: http://www.seg3d.org (2016)

[58] Berron, D., Vieweg, P., Hochkeppler, A., Pluta, J.B., Ding, S.L., Maass, A., Luther, A., Xie, L., Das, S.R., Wolk, D.A., Wolbers, T., Yushkevich, P.A., Düzel, E., Wisse, L.E.M.: A protocol for manual segmentation of medial temporal lobe subregions in 7 Tesla MRI. NeuroImage: Clinical 15(May), 466–482 (2017). https://doi.org/10.1016/j.nicl.2017.05.022

[59] Wisse, L.E.M., Gerritsen, L., Zwanenburg, J.J.M., Kuijf, H.J., Luijten, P.R., Biessels, G.J., Geerlings, M.I.: Subfields of the hippocampal formation at 7t mri: In vivo volumetric assessment. NeuroImage 61(4), 1043–1049 (2012). https://doi.org/10.1016/j.neuroimage.2012.03.023

[60] Yushkevich, P., Avants, B., Pluta, J., Das, S., Minkoff, D., Mechanichamilton, D., Glynn, S., Pickup, S., Liu, W., Gee, J.: A high-resolution computational atlas of the human hippocampus from postmortem magnetic resonance imaging at 9.4 t. NeuroImage 44(2), 385–398 (2009). https://doi.org/10.1016/j.neuroimage.2008.08.042

[61] Ding, S.-L.: Comparative anatomy of the prosubiculum, subiculum, presubiculum, postsubiculum, and parasubiculum in human, monkey, and rodent. Journal of Comparative Neurology 521(18), 4145–4162 (2013) https://arxiv.org/abs/ https://onlinelibrary.wiley.com/doi/pdf/10.1002/cne.23416. https://doi.org/10.1002/cne.23416

[62] Ding, S.-L., Hoesen, G.: Organization and detailed parcellation of human hippocampal head and body regions based on a combined analysis of cyto- and chemoarchitecture. J Comp Neurol 523, 2233–2253 (2015). https://doi.org/10.1002/cne.23786

[63] Insausti, R., Tuñón, T., Sobreviela, T., Insausti, A.M., Gonzalo, L.M.: The human entorhinal cortex: A cytoarchitectonic analysis. Journal of Comparative Neurology 355(2), 171–198 (1995) https://arxiv.org/abs/ https://onlinelibrary.wiley.com/doi/pdf/10.1002/cne.903550203. https://doi.org/10.1002/cne.903550203

[64] Insausti, R., Córcoles-Parada, M., Ubero, M.M., Rodado, A., Insausti, A.M., Muñoz-López, M.: Cytoarchitectonic areas of the Gyrus ambiens in the human brain. Frontiers in Neuroanatomy 13 (2019). https://doi.org/10.3389/fnana.2019.00021

[65] Olga, K., Zilles, K., Palomero-Gallagher, N., Schleicher, A., Mohlberg, H., Bludau, S., Amunts, K.: Receptor-driven, multimodal mapping of the human amygdala. Brain Structure and Function 223 (2018). https://doi.org/10.1007/s00429-017-1577-x

[66] Amunts, K., Olga, K., Kindler, M., Pieperhoff, P., Mohlberg, H., Shah, N., Habel, U., Schneider, F., Zilles, K.: Cytoarchitectonic mapping of the human amygdala, hippocampal region and entorhinal cortex: intersubject variability and probability maps. Anatomy and embryology 210, 343–52 (2006). https://doi.org/10.1007/s00429-005-0025-5

[67] Mallat, S.: Recursive interferometric representations. In: European Signal Processing Conference, pp. 716–720 (2010)

[68] Sifre, L., Mallat, S.: Rigid-Motion Scattering for Texture Classification (2014)

